# Multiphasic myelination and dendritic growth modulate qMRI signals in human visual cortex

**DOI:** 10.64898/2026.05.03.722280

**Authors:** Clara María Bacmeister, Vaidehi S. Natu, Alex Rezai, Sarah Tung, Christina Tyagi, Xiaoqian Yan, Kei Braun, Michael Mui, Congyu Liao, Nan Wang, Xiaozhi Cao, Hua Wu, Patrik Johansson, Annika Enejder, Alexander Beckett, Erica Walker, David Feinberg, Kawin Setsompop, Mercedes Paredes, Kalanit Grill-Spector

## Abstract

How does myelin develop in human visual cortex? By combining immunohistochemistry with *in vivo* and postmortem magnetic resonance imaging of longitudinal relaxation rate (R_1_), which increases with myelin content, we find that myelin and R_1_ increase across development but follow distinct trajectories. Immunohistochemistry reveals two phases of myelination: an infant phase of limited oligodendrogenesis, with myelin restricted to deep cortical layers, followed by widespread myelination across all layers during childhood. Cortical R_1_ also increases across development and correlates with myelin by childhood. However, in infancy, R_1_ increases outpace myelin growth and instead tracks dendritic arborization, indicating that the microstructural drivers of R_1_ change across development. We hypothesize that deep layer myelination in infancy contributes to early visual function whereas later myelination of superficial layers enables prolonged cortical plasticity and learning of complex visual behaviors.

## Main Text

A key question in neuroscience is understanding how cortical tissue develops to support functional development. While classic neuroanatomical research examined the development of neural connections and synapses (*1–3*) or long range white matter connections (*4–6*), more recent research in animals has suggested that myelin–the fatty sheath insulating axons–plays a vital role in shaping cortical circuits. Cortical myelination is modulated by neural activity and experience (*7–18*), influences circuit function (*19–24*), contributes to critical period closure (*25–28*), and, when disrupted, can cause visual and cognitive deficits (*29–36*). Thus, elucidating the development trajectory of cortical myelination in humans is important for determining windows of plasticity and maturation during cortical development. As such, human research has used non-invasive quantitative magnetic resonance imaging (qMRI) of tissue relaxation rate (R_1_[s^-1^]) to glean insights into myelin development (*37–54*). However, qMRI metrics are indirect, impacted by multiple aspects of the cortical microstructure, not just myelin (*55–58*), and these links may be different in the immature, developing cortex. Thus, how human cortex myelinates across development is unknown.

Using human visual cortex as a model system and immunohistochemistry (IHC) to label myelin and oligodendrocyte lineage cells in rare pediatric samples, we first elucidate the developmental trajectory of oligodendrogenesis and myelination in human visual cortex from infancy to young adulthood. Then, we use *in vivo* qMRI of tissue relaxation rate, R_1_[s^-1^]–linearly related to myelin in adults (*56, 57*)–in 121 participants from 0-28 years of age and pair this with postmortem qMRI and IHC to directly test the relationship between myelin and R_1_ in the developing visual cortex.

We use visual cortex as a model system because it is well understood, undergoes profound development in infancy and childhood, and contains regions with different function and developmental trajectories (*42, 59–68*) that are located in consistent anatomical locations that can be reliably identified in postmortem samples (*42, 59–64, 66–71*). Primary visual cortex (V1) is located in the calcarine sulcus (*72–75*) and supports basic visual abilities, such as contrast sensitivity, which develop in infancy (*76, 77*). Higher-level face-selective cortex in the fusiform gyrus (*42, 59–67, 69, 70*) and place-selective cortex in the collateral sulcus (*71*) support face and place recognition, respectively (*59, 78, 79*), and continue to develop during childhood (*42, 80, 81*).

### Sparse, region-dependent myelination of human visual cortex in infancy

*What is the trajectory of myelin development in human visual cortex?* We quantify the developmental trajectory of cortical myelination using IHC for myelin basic protein (MBP, a marker of mature myelin sheaths) in nine rare postmortem pediatric samples of the visual system spanning 0-25 years: six infants (0-6 months), two children (2 and 9 years), and one adult (25 years) (**Fig. 1A, Supplemental Fig. S1**). We measure the percent cortical area covered by MBP in calcarine (Calc, V1), fusiform gyrus (FG, face-selective), and collateral sulcus (CoS, place-selective) in an automated and observer-independent manner (**Supplemental Fig. S1,S2**) and identify cortical layers and the gray/white matter boundary using DAPI, which labels all cell nuclei (**Supplemental Fig. S1,S2**). We test if the development trajectory of myelin coverage (i) occurs early in infancy, (ii) is graded and prolonged into childhood and adulthood, or (iii) is nonmonotonic (e.g. initially rising, peaking, and later declining, **Supplemental Fig. S3**) and if this trajectory varies across cortical regions. One possibility is that the visual system myelinates together to scaffold the later emerging cortical function. Alternatively, myelination trajectories may mirror functional development, predicting that regions which mature earlier functionally, like Calc (V1), may myelinate earlier.

**Fig. 1.**
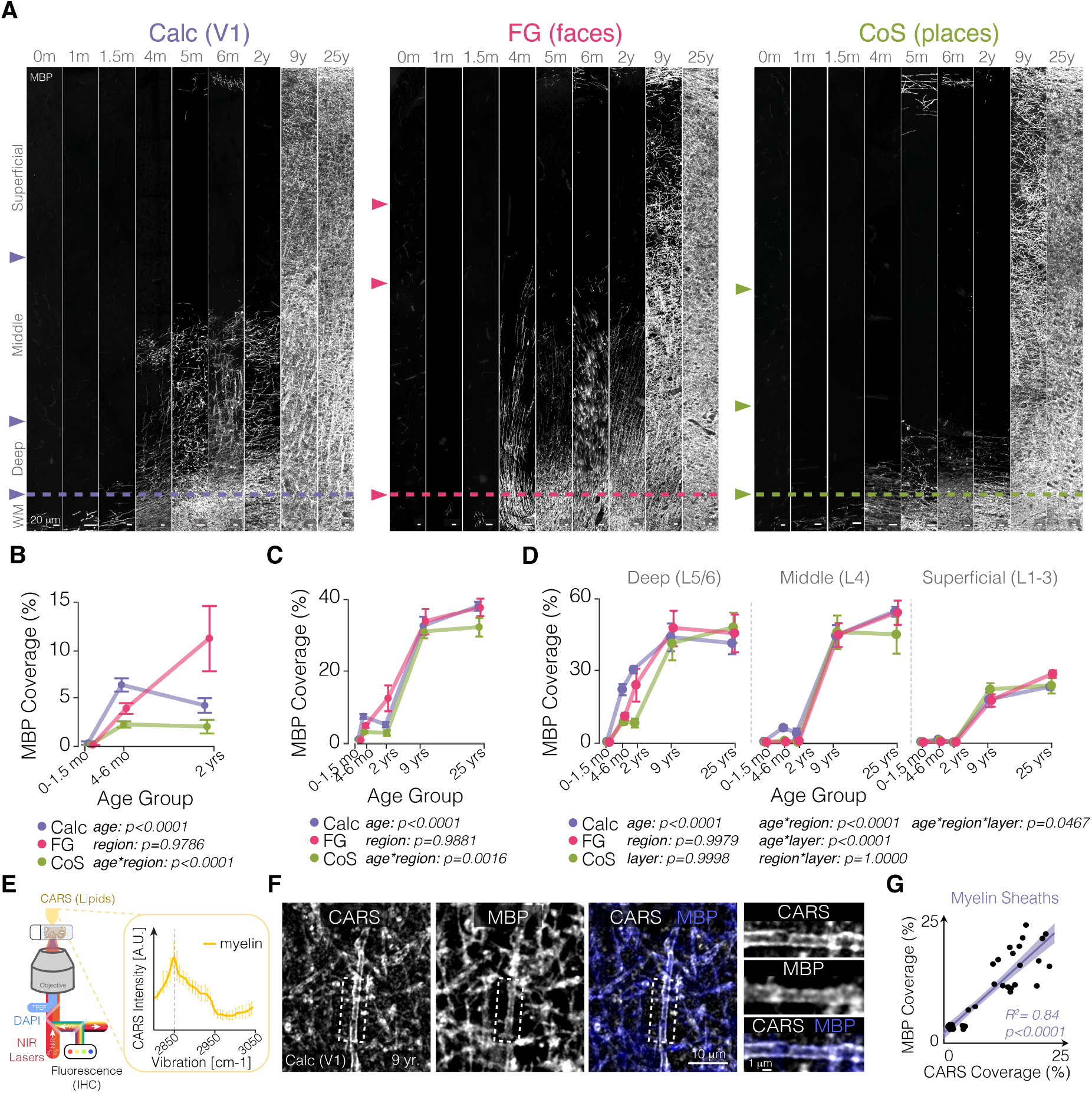
Multiple, region-specific phases of myelination in human visual cortex. **A** Maximum projection of MBP immunostaining across calcarine sulcus (Calc), fusiform gyrus (FG,), and collateral sulcus (CoS) from 0-25 years. Age is indicated on top; *mo:* months; *y:* years. *Horizontal dashed line*: gray matter (GM)–white matter (WM) boundary. Laminar organization is determined from DAPI and varies across regions. **B** Development of myelin coverage in infants (0-2 years) in Calc, (purple), FG (pink), and CoS (green). **C** Same as **(B)** D from 0 to 25 years (0-2-y are same data as **B**). **D** Development of myelin coverage in deep (L5/6, left), middle (L4), and superficial (L1-3) visual cortex. **B-D**: *Dots:* mean; *error bars:* SEM across sections. **E** High-resolution multimodal microscopy, including confocal scanning fluorescence of stained features and label-free Coherent anti-Stokes Raman scattering (CARS) of molecular vibrations emphasizing lipid features, including myelin (2850 cm^-1^, inlay). **F** *Left:* Single-plane image of label-free CARS imaging of lipids, *middle:* MBP-immunostaining of the same section. *right:* overlaid CARS (white) and MBP-immunolabeling (blue) in L6 of Calc in a 9-year-old. *White dotted boxes:* location of magnified inlay on the right. **G**, Positive correlation between compact, mature myelin sheaths labeled with CARS and MBP in Calc; line and shaded area represents line of fit and confidence interval and dots are individual ROIs across depth in 1 month, 6 months, and 9 year old samples.

In all three neonatal samples, visual cortex is devoid of myelin before increasing in 4–6-month-olds (main effect of age: F_2,61_=23.1, p<0.0001, linear mixed model (LMM) with section as random effect); **Fig. 1A,B**). In infancy, myelin development also differs significantly by region (age*region interaction: F_4,61_=10.1, p<0.0001; **Fig. 1A,B**): Calc shows early increases that plateau by 2 years; FG increased throughout infancy, surpassing other regions by 2 years; and CoS showed minimal myelin coverage even at 2 years of age. Extending these measurements to later in development (9 and 25 years), we find that myelin increases substantially during childhood in all three visual regions and into adulthood in Calc and FG (age effect: F_4,81_=140.45, p<0.0001; age*region interaction: F_8,81_=3.51, p=0.0016; **Fig. 1A,C**). These data reveal two phases of cortical myelination: a modest, region-specific increase in infancy followed by widespread development during childhood.

### Myelination starts in deep cortical layers and is prolonged in superficial layers

Myelin coverage is nonuniform across cortical depth (**Fig. 1A**), which may be related to differences in cell types and circuitry between cortical layers (*82*). Quantifying laminar myelin coverage reveals layer-dependent myelin development (age*layer interaction: F_8,249_=13.13, p<0.0001, LMM, section:random effect; **Fig. 1A,D**). During infancy, myelin is largely restricted to deep layers (L5/6) and in Calc extends to layers 4b and 4c (**Fig. 1A,D, Supplemental Fig. S5**). In contrast, during childhood (from 2 to 9 years), myelin coverage increases across cortical layers, with continued development of superficial and middle layers into adulthood (**Fig. 1A,D**).

Laminar patterns of myelination also develop differently by region (age*region*layer interaction: F_16,249_=1.70, p=0.047; **Fig. 1A,D**), reflecting rapid deep-layer myelination in Calc during infancy, limited myelination of CoS (which only occurs between 2-9 years of age), and continued myelination of superficial FG into adulthood. In addition, the orientation of deep-layer myelin differs across visual regions with predominantly tangential sheaths in Calc and CoS and radial sheaths in FG (age*region interaction: F_4,28_=7.087, p=0.0005; **Supplemental Fig. S6**). These data reveal region and layer specific trajectories of cortical myelination.

To independently validate the lack of myelin identified with MBP immunostaining, we use coherent anti-stokes Raman scattering (CARS), a technique that enables label-free imaging of myelin-associated lipids using the unique molecular vibrations of their CH_2_ bonds, in the same sections (**Fig. 1E,F, Supplemental Fig. S7**). Coverage of compact myelin shows strong correspondence across MBP immunostaining and CARS (**Fig. 1F,G)**. That is, when myelin sheaths are present, they are detected by both methods, including in the superficial white matter of neonates (**Supplemental Fig. S7**), and when myelin is absent, it is absent in both measures, providing independent evidence that visual cortex is unmyelinated in neonates.

### Limited differentiation of OL lineage cells in neonates

*Why is cortex sparsely myelinated until childhood?* Myelin is generated and integrated into neuronal circuits by oligodendrocytes, which are produced through a process called oligodendrogenesis. During oligodendrogenesis, oligodendrocyte precursor cells (OPCs) differentiate into premyelinating oligodendrocytes (preOLs), which then mature into myelinating oligodendrocytes (OLs) (**Fig. 2A**). However, 90% of preOLs die before maturing into myelinating oligodendrocytes (*17*). The new OLs that are generated extend and wrap their membranes around axons to form nascent myelin sheaths, which are subsequently compacted into mature myelin via MBP. Therefore, the absence of MBP labeling in the neonatal visual cortex could reflect incomplete oligodendrogenesis, or myelin that is not yet compacted, rendering it undetectable with MBP. To distinguish between these alternatives, we immunostained for Olig2, a marker of oligodendrocyte lineage cells (OPCs, preOLs, and OLs), and BCAS1, a marker of preOLs and new myelinating OLs that is expressed before MBP (*75*) (**Fig. 2A-D**). We reasoned that Olig2+ cells would reflect overall oligodendrocyte lineage cells, while BCAS1+ stained cells would indicate differentiation, reflecting preOLs, new OLs, and new myelin. Olig2+ cells are present in neonates across cortical regions (significantly > 0, p<0.0001 for all regions and layers, one sample t-test), and their density does not differ with age in infancy (main effect of age: F_2,5039_=0.14, p=0.71; LMM, section:random effect, **Fig. 2C,E, Supplemental Fig. S8**). In contrast, across visual cortex, BCAS1+ cells were low in neonates (main effect of age: F_2,39_=5.15, p=0.0104, **Fig. 2D,F**), and MBP+ sheaths are absent (**Fig. 1A-C**), indicating little differentiation of oligodendrocyte lineage cells in neonates. Corroborating this, the superficial white matter of Calc contains both BCAS1+ cells and myelin at birth (**Supplemental Fig. S9**), confirming that the absence of cortical myelination and differentiating OLs is not due to lack of staining sensitivity in infants. By 4-6-months, BCAS1+ cell density increases markedly in the cortex and then declines by ∼50% at 2 years (**Fig. 2F, Supplemental Fig. S8**), highlighting a burst of differentiation that parallels the onset of cortical myelination.

**Fig. 2.**
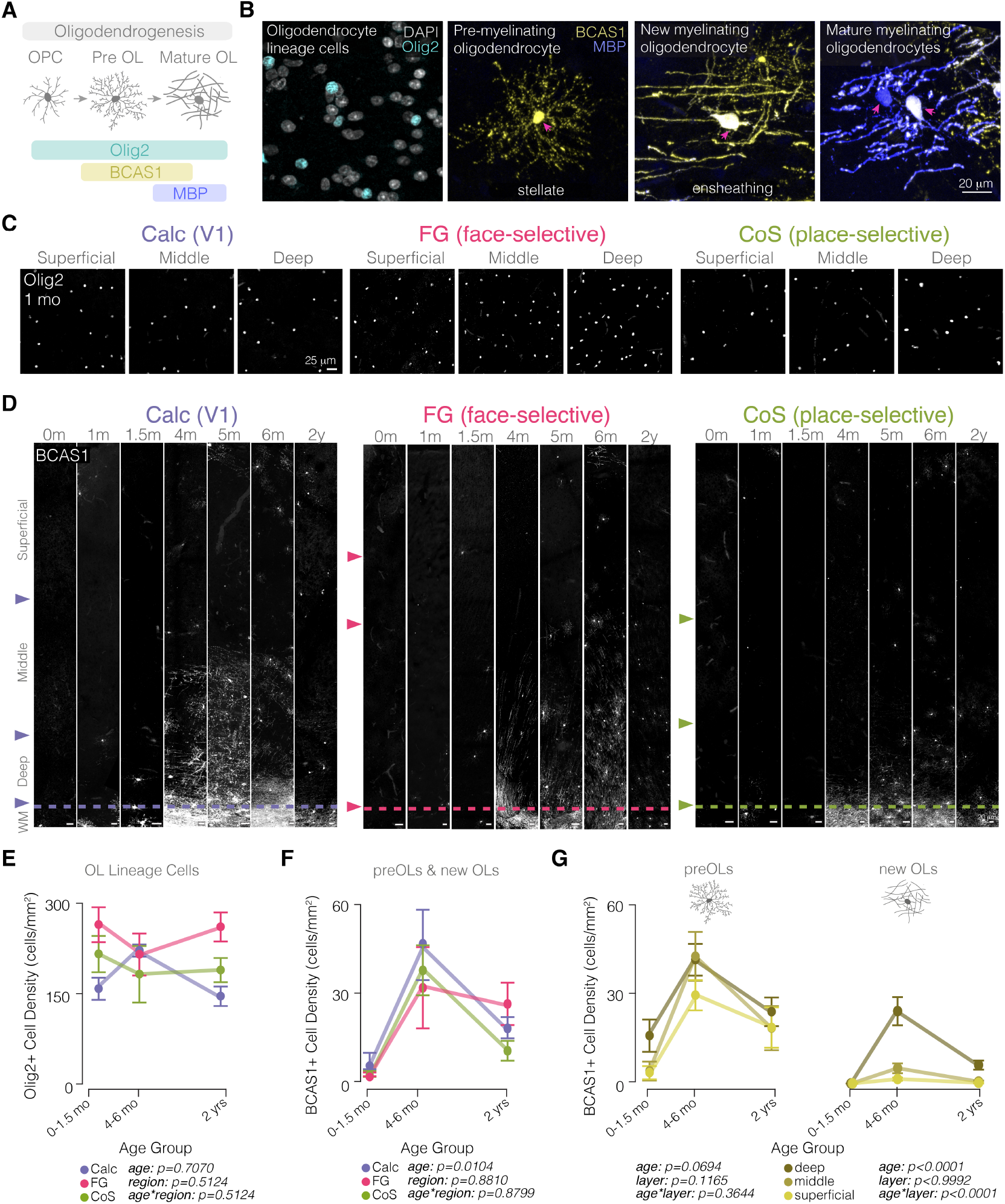
Age-dependent progression of oligodendrogenesis in developing visual cortex. **A** Schematic of oligodendrogenesis. **B** Maximum projection of Olig2 (cyan), BCAS1 (yellow) and MBP (blue) immunostaining showing three cellular stages: BCAS1+ stellate preOLs, BCAS1+ ensheathing new OLs with limited MBP+ staining, and MBP+ mature OLs with reduced BCAS1+ staining. Pink arrows point to cell bodies. **C** Maximum projection of Olig2 in neonate (1-mo) Calc (purple), FG (pink), and CoS (green) in superficial, middle, and deep cortical layers. **D** Maximum projection of BCAS1 immunostaining across Calc (purple), FG (pink), and CoS (green) from 0-2 years of age. Age indicated on top, m: months, y: years. **E,F**, Mean Olig2+ (**E**) and BCAS1+ (**F**) cell density across Calc (purple), FG (pink), and CoS (green) from 0-2 years of age. **G**, Mean density of preOLs (left, stellate) and new OLs (right, ensheathing) across deep (dark yellow), middle (yellow), and superficial (light yellow) cortical layers from 0-2 years. Error bars: SEM across sections.

### BCAS1+ new OL generation is restricted to deep cortical layers in infancy

We next asked whether layer-specific myelination in infancy is related to laminar distribution of differentiating OL lineage cells. BCAS1+ preOLs are evenly distributed across layers (**Fig. 2D,H**, no main effect of layer: F_2,132_=2.19, p=0.11654) and develop similarly across layers (no interaction effect of age*layer: F_4,132_=1.09, p=0.3644). In contrast, BCAS1+ newly-formed OLs increase in a layer-dependent manner (interaction effect of age*layer: F_4,72_=13.06, p<0.0001; **Fig. 2H**), with significantly higher densities of BCAS1+ new OLs in deep layers compared to superficial and middle layers of cortex in 4-6-month- and 2-year-olds (post-hoc t-test, p<0.001 for all comparisons). This deep layer enrichment of new myelinating OLs mirrors the laminar pattern of early myelin accumulation: minimal myelination in neonates mirrors limited OPC differentiation, while spatially restricted myelination in infants mirrors the deep-layer bias of new OLs in infants.

### Development of R_1_ outpaces development of myelin coverage in infant visual cortex

*Can cortical R*_*1*_ *be used as a biomarker to track cortical myelination in developing children?* To answer this question, we obtained qMRI of R_1_ in 121 individuals spanning 0-27 years of age (27/45 infants ages 0-15-months provided longitudinal data). Cortical R_1_ shows profound development during infancy that extends into childhood (**Fig. 3A, Supplemental Fig. S2**). We quantified R_1_ development by projecting V1, mFus-faces, and CoS-places from adult atlases (*83*) into each individual’s brain and measuring R_1_.

**Fig. 3.**
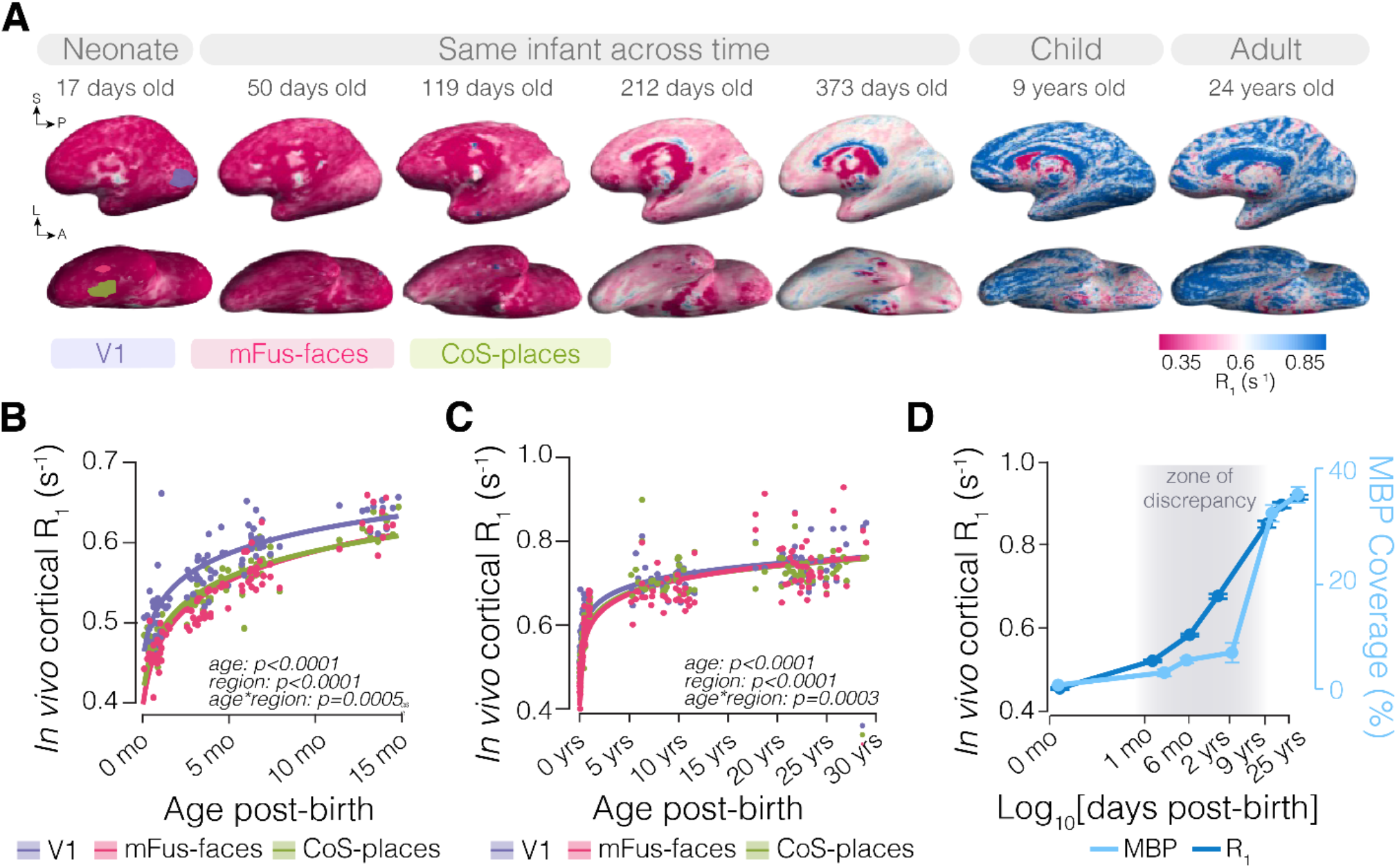
A wave of myelin-independent R_1_ development in infant visual cortex. **A**, R_1_ in an example neonate (17-days-old), an example infant across year one year (at 50, 119, 212, and 373 days), and a 9-year-old and 24-year-old on their inflated right-hemisphere cortical surfaces. Atlas V1 (purple), mFus-faces (pink), and CoS-places (green) projected on the example neonatal cortical surface. **B**, R_1_ development in V1 (purple), mFus-faces (pink), and CoS-places (green) from 0 to 15 months. **C**, same as (**B**) from 0 to 28 years (0-15 months are the same as in **B**). Each dot is data from an individual scan colored by region. **D**, R_1_ (dark blue) and MBP coverage (light blue) averaged across regions from 0 to 25 years of age. Dots and error bars: mean and SEM across scans for R_1_ and sections for MBP.

In infancy, R_1_ increases faster from 0-6 months than 6-12 months, with differential development across regions (age*region: F_2,254_=5.24, p=0.0059, LMM with random variable of participant, **Fig. 3B**). At birth, R_1_ was highest in V1 and during the first year of life, mFus-faces developed at a faster rate than V1 and CoS-places (**Supplemental Fig. S10**). These differences persisted across the 0-27 age range (age*region: F_2,482_=20.94, p<0.0001, **Fig. 3C**), with prolonged maturation of mFus-faces relative to V1 and CoS-places (**Supplemental Fig. S10**). Thus, cortical R_1_ exhibits rapid growth during infancy that continues during childhood with region-specific developmental trajectories.

We next sought to compare cortical R_1_ and cortical myelin coverage across the three visual regions of interest (**Fig. 3D**). We reasoned that although these metrics were measured in different samples and resolutions, if cortical R_1_ is driven by myelin content, these metrics should have a similar developmental trajectory. While both R_1_ and myelin are lowest in neonates and highest in adults, we find a zone of discrepancy in infancy. Myelin coverage increases modestly between 0-2 years, reaching only a quarter of its growth by age 2 then almost quadrupling between 2 and 9 years of age (**Fig. 3D, Supplemental Fig. S11**). In contrast, R_1_ rises rapidly during the first year of life, reaching half of its total growth by ∼1 year of age and then continuing to increase during childhood. This divergence reveals a dissociation between R_1_ and myelin development in infancy.

### Myelin predicts cortical R_1_ in childhood, but dendrites modulate R_1_ in infancy

*Given that infant visual cortex is sparsely myelinated, what drives the pronounced increase in cortical R*_*1*_ *during infancy?* One possibility is that, in the absence of myelin, other aspects of cortical microstructure may grow and increase R_1_ in infancy, e.g. dendritic arborization continues in V1 postnatally (*84–88*), which may increase tissue density and consequently R_1_. Alternatively, the comparison between *in vivo* (R_1_) and postmortem (MBP) metrics in different samples and resolutions may contribute to the observed divergence. To distinguish between these possibilities, we directly assess the relationship between R_1_ vs. cortical myelin (MBP coverage) and dendritic arborization (MAP2, a marker of neuronal dendrite coverage, **Fig. 4A,D, Supplemental Fig. S12**) in the same regions in 4 postmortem samples spanning development (1.5 months, 5 months, 9 years, and 25 years old, **Fig. 4A**). Postmortem R_1_ was obtained at 0.5mm resolution, compared to 1-2mm *in vivo* resolution enabling multiple measurements of R_1_, MBP coverage, and dendritic coverage across cortical depths.

**Fig. 4.**
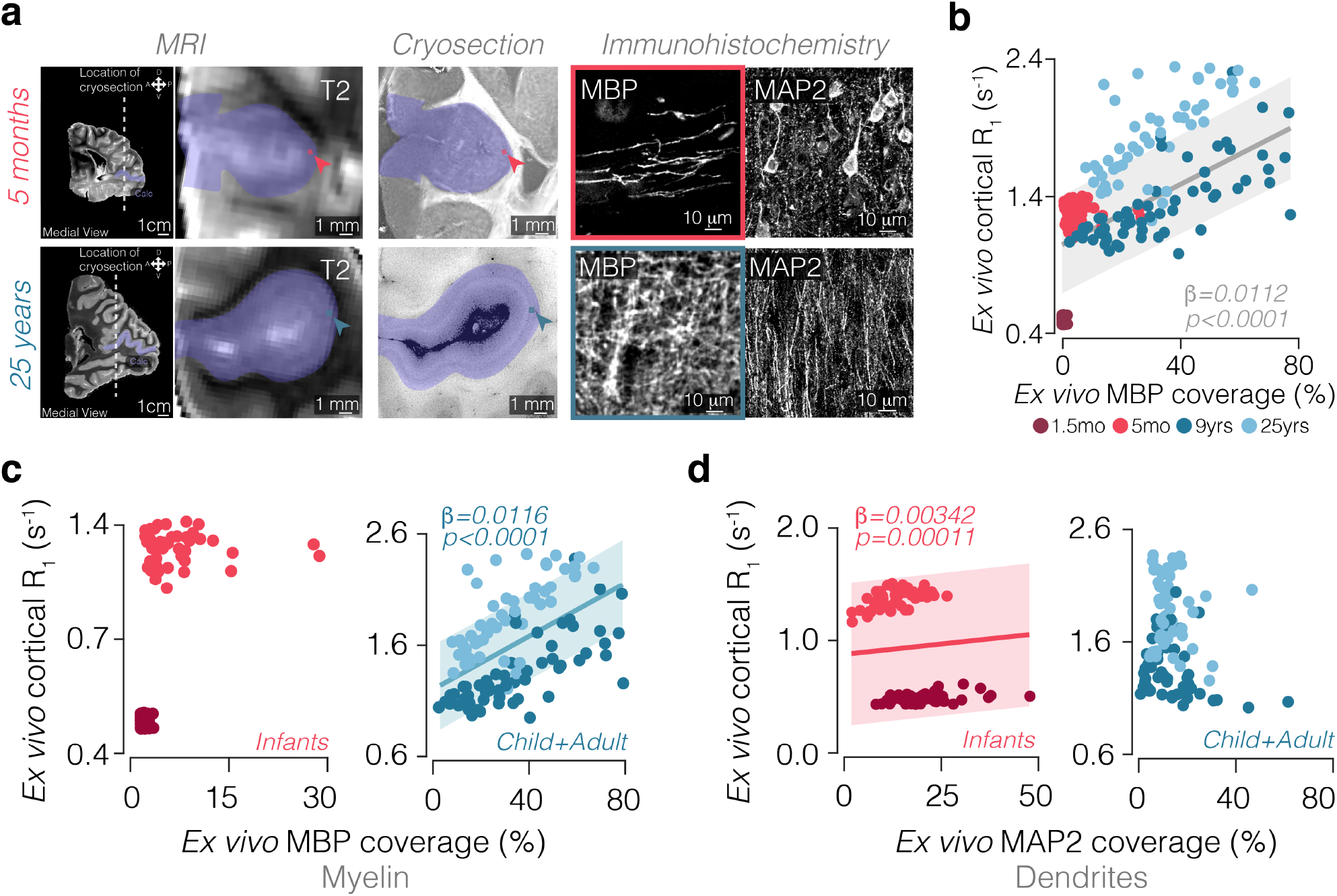
R_1_ tracks dendritic arborization in infancy and myelination later in development. **A** Tissue alignment from postmortem MRI of postmortem samples (two left panels) to cryosections (middle) to immunohistochemistry of an ROI imaged on a confocal microscope (right, MBP) and the adjacent section for MAP2. Top: 5-months-old, Calc; Bottom: 25-year-old, Calc. Strawberry and teal squares highlight the same ROI in postmortem MRI, cryosections, and confocal images of IHC, immunostained for myelin (MBP) and dendrites (MAP2). **B** Relation between R_1_ and MBP coverage in visual cortex of the same postmortem samples. **C** R_1_ as a function of MBP coverage in infant (*left*) and child and adult samples (*right*). **D** R_1_ as a function of MAP2 coverage in infant (*left*) and child and adult samples (*right*). **B-D** *dark red:* 1.5-month-old, *strawberry*: 5-month-old, *dark teal*: 9-year-old, *light teal:* 25-year-old. *Dots:* individual ROIs across cortical depth.

In general, R_1_ is linearly related to myelin coverage (**Fig. 4B**, beta=0.0012, p<0.0001, linear mixed model (LMM), sample is a random intercept). However, the neonate lies completely outside of the confidence interval of this regression, suggesting that myelin significantly contributes to R_1_ when present, but that other tissue features may contribute to cortical R1 in the absence of myelin, as in infant cortex. To test this prediction, we split our samples into sparsely-myelinated infant samples (1.5 and 5 months) and myelinated child and adult (9 and 25 years) samples (**Fig. 4C**) and chose regions that that maximized the range of myelination (i.e. Calc/V1 is most myelinated and CoS is the least in infancy) and have similar anatomical structure, i.e. both are sulci (**Fig. 3A**). Indeed, in child and adult samples, R_1_ linearly increases with MBP coverage (beta=0.016, p<0.0001, LMM, sample random intercept). However, in infant cortex, there is no significant relationship between R_1_ and myelin (beta=0.000226, p=0.88, LMM). In contrast to MBP coverage, MAP2 coverage increases rapidly in the first two years of life in all visual regions and then stabilizes (main effect of age: F_5,29_=7.40, p=0.0002, age*region interaction: F_10,42_=1.33, p=0.25; LMM, section-random effect, **Supplemental Fig. S12**). Strikingly, in infants, R_1_ significantly and linearly increased with MAP2 coverage (infants: beta=0.0034, p=0.00011, LMM). In contrast, R_1_ does not significantly increase with MAP2 in child and adult samples (beta=-0.0442, p=0.16, LMM, **Fig. 4D**). As the contribution of MAP2 in infants (beta=0.0034) is more modest than MBP (beta=0.0116), it may be masked later in development and only visible in sparsely myelinated cortex. Together, these results suggest that R_1_ cannot be used solely as a biomarker for cortical myelin development. Instead, we find that dendritic arborization contributes to increases in cortical R_1_ in the first year of life, while myelin becomes the primary contributor by late childhood.

## Discussion

We find that human visual cortex develops through multiple stages of microstructural growth, each predominated by distinct cellular processes: dendritic arborization in infancy and myelination in childhood. In neonates, both primary and high-level visual cortex are unmyelinated, and visual cortex undergoes profound dendritic arborization. By 4 to 6 months, myelination begins, yet it remains sparse and confined to deep cortical layers. Premyelinating and new oligodendrocytes differentiate in the same developmental period and layers, respectively, in which myelination begins, suggesting that distinct stages of oligodendrogenesis determine the laminar and temporal patterning of cortical myelination in early development. In childhood, there is widespread myelination across all cortical layers, with particularly protracted myelination of the superficial layers of the FG.

We hypothesize that rapid dendritic arborization across visual cortex in infancy may scaffold synaptic connectivity, and myelination of deep layers in infancy may stabilize long-range connections and enhance efficient signal transmission (*15, 25, 28, 89*), together enabling early visual function. In contrast, the sparse myelination of superficial layers containing cortico-cortical connections may support prolonged cortical plasticity (*11, 16, 17, 26–28*) (*33–35*), enabling learning of complex visual behaviors like face recognition and navigation through childhood and adolescence (*90*). It is interesting that the trajectory of myelin development and oligodendrogenesis parallels functional development. Regions that show earlier visual activity, such as V1 and face-selective cortex (4-6-months (*59, 79, 91*)), start myelinating before later emerging place-selective cortex (*79*), and regions with a protracted functional development through adolescence, such as face-selective cortex in the FG (*61, 63, 66, 68*), show the most extended superficial-layer myelination.

Ultimately, it is desirable to directly link cellular and functional development within the same infant over the course of development, necessitating *in vivo* neuroimaging measurements of microstructure and function. We find that the cellular basis of cortical qMRI signals changes across development: dendritic proliferation is a dominant factor modulating cortical R_1_ in infants, while myelin predominates cortical R_1_ by childhood. These data caution against using cortical R_1_ as a biomarker for myelin in early development (*37–54*) when cortex is immature and sparsely-myelinated but validate its use for assessing myelination in childhood and young adulthood. Future studies can test additional aspects of cortical cellular development such as axonal branching (*92*), glial development (*93*), and iron content (*94*) and evaluate their impact on R_1_ (*55–58*) and additional qMRI metrics such as transverse relaxation rate 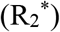, proton density (PD) (*55, 57*), and relaxivity (*95*) to provide further insights of how multimetric qMRI may be used to track separate aspects of microstructural development in cortex. Together, our study enhances understanding of cortical myelination across development and offers a roadmap for interpretable *in vivo* diagnostic tools for early detection of neurodevelopmental atypicalities.

## Acknowledgments

We thank Stanford Cell Science Imaging Facility (SCIF) for providing microscopy equipment. We thank Drs. Carla Shatz, Brad Zuchero, Juliet Knowles for fruitful conversations and feedback on the study.

## Author contributions

CMB conceived the project, designed and performed experiments, analyzed data, generated all figures, and wrote the manuscript with input from other authors.

VSN designed experiments and collected and analyzed data for in vivo and postmortem scans.

AR performed experiments and analyzed data for Fig. 2 and Supplemental Fig. S8. ST, CT, and XY collected data for in vivo infant scans.

KB analyzed data for Supplemental Fig. S6.

MM performed tissue processing and provided technical support.

CL, NW, XC collected data and generated sequences and reconstructions for postmortem scans at 3 Tesla at CNI and 7 Tesla at Berkeley.

HW designed sequences for in vivo infant scans.

AE obtained funding for the project, developed multi-modal CARS/fluorescence microscopy and analysis for myelin imaging, designed and contributed the CARS/fluorescence experiments, data analysis, and writing.

PJ developed and designed the multi-modal CARS/fluorescence microscopy experiments and analyzed the CARS/fluorescence data.

AB, EW, and DF contributed sequences and scans for the postmortem infant at 7 Tesla at Berkeley.

KS designed sequences and reconstructions for postmortem scans at 3 Tesla at CNI and 7 Tesla at Berkeley.

MFP conceived of and obtained funding for the project, obtained postmortem samples, supported the IHC data collection and analyses and writing of the manuscript.

KGS conceived of, designed, obtained funding and oversaw the project, contributed to the data analysis and wrote the manuscript with input from other authors.

## Data, code, and materials availability

All data, code, and materials used in the analysis are available on Github: https://github.com/VPNL/bbmyelin.

## Supplementary Materials

Materials and Methods

Supplementary Text

Figs. S1 to S12

Tables S1 to S4

References (*1-100*)

