## Supplemental Figures and Tables for "Multiphasic myelination and dendritic growth modulate qMRI signals in human visual cortex"

**The PDF file includes:**

Materials and Methods
Figs. S1 to S12
Tables S1 to S4

### Materials and Methods

#### Postmortem samples

***Sample preparation.*** Pediatric and adult brain samples were obtained via the UCSF Pediatric Neuropathology Brain Bank. Nine brain samples include six infants: newborn, 1, 1.5, 4, 5, 6 months; two children: 2, 9-year-old, and one adult (25-year-old). The 1.5-months, 5-months, 9-year-old, 25-year-old samples included the entire occipital lobe and the posterior half of the temporal lobe and underwent MRI prior to cryosectioning and staining (**Supplemental Fig. S1**). Samples were obtained from clinical cases without neurological medical history with a postmortem interval of 24-48 hours (see **Supplementary Table 1**). Brains were postfixed in 0.5% PFA for long-term storage at 4 °C, transferred to graded solutions of 10%, 20%, and finally 30% sucrose solution in PBS (pH 7.4), and stored at 4 °C for at least 24 h at each step. Brains were then sectioned into blocks containing our three regions of interest, frozen in TissuePlus O.C.T, and sectioned at 30 μm thick.

***Identification of regions for analysis.*** Sulcal and gyral landmarks were identified on the intact postmortem hemispheres to guide anatomical registration (*68*, *69*, *71*, *73*, *81*). We focused on three human visual areas spanning the visual ventral stream processing hierarchy (*75*, *81*): primary visual cortex (V1, in the calcarine sulcus (*72–75*)), high-level face-selective cortex (in the fusiform gyrus, FG(*69*)) and place-selective cortex (in the collateral sulcus, CoS (*71*); **Supplemental Fig. 2**). Each hemisphere was then manually dissected into cortical blocks corresponding to target regions of interest and then cryosectioned at 30 μm thickness. Sulcal and gyral features were re-identified at each stage of processing to ensure anatomical continuity using stereotypical anatomical locations: V1 is in the calcarine sulcus (Calc) and possesses the stria of Gennari; face-selective regions overlap the mid fusiform sulcus and extend to the lateral fusiform gyrus, which has a distinct ω-shape in coronal sections (*69*); and the most probable location of place-selectivity (Cos-places/PPA) is the intersection between the anterior lingual sulcus (ALS) and the collateral sulcus (CoS, (*71*)) (**Supplemental Fig. S2**). To facilitate precise localization of cortical regions, serial cryosections were immunostained with NeuN and/or DAPI at regular 1 mm intervals throughout each block. These stains were used to identify cytoarchitectonic features and stereotyped laminar patterns that demarcate cortical regions, as described in prior studies. On each immunostained section, cortical layers and the gray:white boundary were delineated by changes in cell density as measured by DAPI staining of nuclei (**Supplemental Fig. S2,3**)

#### Tissue processing and microscopy.

***Immunohistochemistry.*** Sections were acquired at 1mm intervals (mean distance apart: 857.72 μm) across our cortical volume of interest and immunostained (for a full list of antibodies, see **Supplemental Table 2**). Immunostaining was performed on slide-mounted cryosections. Sections were pre-incubated in blocking solution (5% normal donkey serum, 2% bovine g-globulin, 0.3% Triton X-100 in PBS, pH 7.4) for 1-4 h at room temperature, then incubated overnight at room temperature in primary antibody (listed along with secondary antibodies in **Supplementary Table 2**). Secondary antibody incubation was performed at room temperature for 2.5 h. Sections were mounted on slides with Vectashield antifade reagent (Vector Laboratories). Images were acquired with a laser-scanning confocal microscope (Leica DMi8 Inverted Microscope with Adaptive Focus Control and an 8kHz Tandem Scanner STELLARIS 8).

***Multimodal microscopy: Confocal fluorescence and label-free imaging of myelin with Coherent anti-Stokes Raman scattering (CARS).*** For CARS imaging of calcarine sulcus (Calc), sections were immunostained as above but were mounted in PBS rather than mounting medium. Mounting media includes glycerol, which contains methylene groups, contributing to resonant signals that interfere with the lipid signature. To allow us to perform spectral evaluations enabling us to distinguish between various lipid features (e.g. lipid droplets vs myelin) characterized by different spectral signatures, we therefore mounted our slides using PBS. Similarly, when sealing slides with traditional methods, such as nail polish, hydrophobic components may leech into the slide and colocalize with features of interest in nonlinear microscopy. Therefore, to prevent samples from drying out, slides were sealed with VaLaP a 1:1:1 mixture of vaseline, lanolin, and paraffin.

A Nikon Ti2-E microscope with a C2 mirror scanner controlled by NIS-Elements Software (Nikon) equipped with an objective with high numerical aperture and magnification (Apo TIRF, NA 1.49, 100X) and customized to include a pico-second laser system (picoEmerald S) with 2 picosecond pulse length and 80 MHz repetition rate was used to acquire all CARS images. The laser system includes a 1031 nm NIR pump laser and an optical parametric oscillator (OPO) with a tunable output between 700-960 nm to generate the CARS signal. CARS was measured in the forward direction, and NIR two-photon excited fluorescence (TPEF) in the epi-direction. The microscope was switched to conventional linear fluorescence with four excitation sources (405, 488, 561, and 640 nm) and tunable detection windows, enabling direct correlation between confocal fluorescence imaging of immunostaining for MBP- and DAPI-labeled cryosections with the signal obtained by CARS microscopy. To get data for lipids, CARS was collected at 2850 $cm^{-1}$ for each high-resolution stack.

First, a low-resolution 20X image (512 x 512, scan rate 1.9 us/pixel dwell time) was acquired to identify cortical regions of interest (ROIs, Calc, Cos, Fus), cortical layers (L1-L6), and the superficial white matter using DAPI profiles, as described above. From this lower-resolution image, 20 evenly spaced, high-resolution fields of view were identified across cortical depth and into superficial white matter for high resolution paired CARS and fluorescence microscopy at 100X magnification. We co-registered CARS with IHC for MBP immunostaining using DAPI in the same sections and identified CARS-emitting cross-sections of myelin sheaths as high-intensity tubular features flanking unlabeled putative axons (**Fig. 1F, Supplemental Fig. S7**).

**Microscope image processing and analysis.**

***Quantification of myelin and dendrite coverage.*** To quantify myelin and dendrite content across cortical layers, custom ImageJ (FIJI, (*96*)) macros were used to segment and analyze MBP and MAP2 immunofluorescence, as well as label-free CARS images of lipids within anatomically defined ROIs. Cortical layer ROIs were manually defined and saved for each image. ROIs were exported using the ROI Manager and stored in a standardized naming format for traceability. For the evaluation of MBP-labeled myelin and dendrite coverage, the image channels corresponding to MBP and MAP2 immunolabeling were duplicated from maximum intensity projections. To reduce noise and correct for background heterogeneity, a Gaussian blur ($\sigma = 1.0$) was applied. In the case of dendrites, a rolling average of 60 was used to reduce noise and correct for background heterogeneity. The processed image was binarized using Otsu's thresholding method (*97*) and analyzed using the Analyze Particles function in ImageJ. All defined layer ROIs were overlaid, and particle analysis was performed within each ROI to quantify the proportion of cortex with immunolabeling (MBP+ or MAP2+). Blood vessels were identified as features that stained across all images and were removed semi-automatically by generating a mask of co-localized staining and then removing it using customized ImageJ macros. Images were then manually checked to ensure all blood vessel features were removed. Additionally, all sections and images were manually reviewed to inspect for folds and tears, which were identified in sections as visible overlaps in tissue and/or rips in the tissue and in images as areas with irregular borders which produced high-intensity signals (folds) or no detectable signal (tears). These areas were excluded from analyses of overall cortical area. Additionally, in infants, neuronal soma were often labeled by MAP2 (see **Fig 4, Supplemental Fig. S12**). To account for this, a semi-automated approach was taken to exclude MAP2-labeled neuronal soma: first, MAP2-labeled sections were also labeled with DAPI/NeuN and, from these channels, a mask was generated to identify and remove DAPI/NeuN and MAP2+ co-localized areas. Then, images were manually inspected to ensure all neuronal soma were segmented and excluded from analysis of MAP2+ dendrites. Summarized data—including region, integrated density, and particle count—were exported for downstream statistical analysis.

***Quantification of CARS coverage.*** Single-plane ROIs (77 $\mu m^2$) spanning cortical depth in child and infant brains were analyzed using a thresholding algorithm (Otsu (*97*)) with a Gaussian blur of $\sigma$ = 5.0 with a rolling average of 20 to reduce noise and correct for background heterogeneity. In cases where CARS coverage was high (>70%), images were manually reviewed for non-myelin sheath related CARS features (see **Supplemental Fig. S7**), which were identified and excluded from analysis to ensure that coverage of only mature myelin sheaths were analyzed. TArea coverage was then corroborated with manual counting of CARS- and MBP-labeled sheaths, which were identified by intense, tubular features in the case of CARS and MBP+ tubular structures in the case of sections stained with MBP (**Supplemental Fig. S7**).

***Myelin sheath orientation analysis.*** Max projections of MBP-immunostained sections were processed with a 1-pixel Gaussian blur filter to aid in the identification of myelin internodes. Individual myelin sheaths were identified by increase in fluorescence intensity for myelin sheaths followed by a decrease to zero in fluorescence intensity for the terminal of the sheath. Individual myelin sheaths were traced using the Simple Neurite Tracer plugin in FIJI (*98*). Tracing was performed on high-resolution confocal image stacks of MBP-labeled cortical sections obtained above. To ensure the robustness and reproducibility of sheath-level reconstructions, two independent observers traced each visible internode manually. Inter-rater reliability was assessed by correlating sheath measurements between observers, yielding a high degree of agreement (Pearson's r > 0.95). Tracings were exported as ROIs for analysis of sheath orientation. Data in **Supplemental Fig. S6.** To assess the orientation of individual myelin sheaths, custom ImageJ (FIJI) macros were used to calculate the overall angle of each trace relative to the horizontal axis. Start and end coordinates of each ROI were extracted to define a direction vector, from which the angle was computed using the arctangent function. All angles were generated within a 0–180° range due to bidirectionality of tracing, and each angle was then normalized to 0–90° range relative to the pial surface.

**Quantitative magnetic resonance imaging (qMRI).**

***Participants.*** We recruited 82 full-term, healthy infants (ages, 0-15 months, 35 females, 47 males), 32 children (5-12 years old, 14 females, 18 males), and 44 adults (ages 22-28, 21 females, 33 males). We obtained usable data from 45 infants (20 females), all children, and all adults. 27/45 infants provided longitudinal data, participating in two or more timepoints (**Supplemental Fig. S1**). The recruitment was carried out in line with racial and ethnic diversity representative of the San Francisco Bay Area (2 Hispanic, 6 Asian, 24 White, and 15 multiracial participants. The procedures were approved by the Stanford University Internal Review Board. All participants or their guardians provided written, informed consent.

***MRI (magnetic resonance imaging) procedure.*** MRI protocols adhered to Stanford University Internal Review Board standards. Scans were scheduled in the evening aligning with the infant's bedtime to minimize discomfort and movement. Upon arrival, caregivers provided written, informed consent. Infants were checked for MRI safety, dressed in MR-safe attire, and swaddled to minimize movement. Noise reduction was achieved using earplugs and neonatal noise attenuators. An MR-compatible plastic immobilizer was used for newborns to stabilize their head position. Throughout the scanning, an infrared camera monitored the infant, and an experimenter remained in the MRI suite to ensure the infant's comfort and promptly address any distress.

***Data quality assurance during MRI.*** Real-time monitoring and post-sequence assessments were crucial. Scans were repeated if excessive head motion or image blurring was detected, with approximately 50% of scans successful on the first try.

***Data acquisition.*** All participants underwent multiple MRI scans during each session to obtain anatomical MRI and quantitative MRI (qMRI). Scanning was performed using two identical 3T GE Discovery MR750 Scanners at Stanford University's Center for Cognitive and Neurobiological Imaging (CNI) and Lucas Imaging Center. Identical acquisition protocols were employed across scanners. For safety, all imaging was conducted at Normal Specific Absorption Rate (SAR) levels due to the infants' low body weight.

***Anatomical and quantitative MRI parameters.*** Subjects were scanned using methods described in our prior work: infants (*45*), children and adults (*42*). Briefly, T1- and T2-weighted images were obtained for tissue segmentation using specific parameters.
*Infants: T1-weighted parameters.* GE 3D BRAVO sequence with TE=2.7 ms; TR=6.7 ms; echo train length=1; voxel size=1 $mm^3$; scan time ~3 min. *T2-weighted parameters:* GE CUBE sequence with TE=122 ms; TR=3650 ms; echo train length=120; voxel size=1 $mm^3$; scan time ~4 min; FOV = 20.5cm. Spoiled gradient echo (SPGR) images were used to generate synthetic T1-weighted images. An inversion-recovery EPI (IR-EPI) sequence with multiple inversion times (TI) was used to estimate $T_1$ (and $R_1$, i.e. $1/T_1$) in each voxel, via slice-shuffling technique: 20 TIs with the first TI=50ms and a TI interval of 150ms, and a second IR-EPI with a reverse-phase encoding direction. For the qMRI sequence for T1, TR=3150ms, TE=47ms, acquisition matrix =100, voxel size = $2mm^3$; number of slices=60, FOV-20cm; in-plane/through-plane acceleration=⅓, with a scan time of 1 min and 45 seconds.
*Children and Adults:* Four spoiled gradient echo images were acquired with different flip angles (4°, 10°, 20°, and 30°), with the following parameters: TE = 2.4 ms; TR = 14 ms; voxel size = 0.8mmx0.8mmx1mm; number of slices = 120; FOV = 22.4 cm, each with a scan time of approximately 4 minutes and 55 seconds. To remove field inhomogeneities, we collected four additional spin echo inversion recovery scans with an echo planar imaging (EPI) readout, a slab inversion pulse, and spectral spatial fat suppression (SEIRs acquired with a TR of 3s, echo time set to minimum full, and 2x acceleration; inversion times 50, 400, 1200, and 2400 ms and were collected

at 2x2mm$^2$ in-plane resolution and slice thickness=4mm. Full scan parameters listed in **Supplementary Table 3**.

***Quantitative T1 relaxation time modeling.***
We analyzed all data in each individual's native brain. All data was aligned to the whole brain 1mm anatomy.
*Infants:* T1 relaxation was modeled using an inversion-recovery signal equation with an exponential decay function with a decay constant $T_1$:

$$(1) \qquad S(t) = a(1 - be^{-t/T1})$$

where t represents the inversion time, a is proportional to the initial magnetization of the voxel, and b is the effective inversion coefficient. Distortion corrections on IR-EPI images were executed using FSL's top-up tool. The Levenberg-Marquardt algorithm (*99*) was applied to fit the corrected images, generating voxel-wise $T_1$ estimates and model fit quality ($R^2$ values).
*Children and Adults:* Synthetic T1-weighted whole-brain images were generated by integrating SPGR and IR-EPI data using mrQ software (https://github.com/mezera/mrQ). We used mrQ (https://github.com/mezera/mrQ) to estimate qMRI parameters and generate synthetic whole brain T1-w anatomies. qMRI were calculated from spoiled-gradient echo images acquired with different flip angles (α = 4°, 10°, 20° and 30°, TR = 14 ms, TE = 2.4 ms) and a voxel resolution of 0.8x0.8x1 mm$^3$, which was resampled to 1 mm$^3$ isotropic. Anatomical data were aligned to the AC-PC plane. From the SPGRs and IR-EPI scans, synthetic T1-weighted whole-brain images were generated

***Generation of cortical surfaces.***
*Infants:* Cortical surfaces were reconstructed using both T2-weighted and synthetic T1-weighted images. Initial segmentations were produced using FreeSurfer's infant-recon-all module (https://surfer.nmr.mgh.harvard.edu/fswiki/infantFS) and further refined with the iBEAT toolbox (Brain Extraction and Analysis Toolbox, iBEAT, v-2.0 cloud processing, https://ibeat.wildapricot.org/). Manual corrections were applied using ITK-SNAP software (http://www.itksnap.org/). The final aligned and corrected segmentations were reinstated into FreeSurfer using custom functions available at (https://github.com/VPNL/babies_graymatter).
*Children and adults:* Synthetic $T_1$ anatomies underwent automated cortical surface reconstruction using FreeSurfer (http://surfer.nmr.mgh.harvard.edu/). The anatomical images were segmented into white and gray matter. White matter surfaces were inspected and manually fixed for missing or mislabeled white matter voxels using ITK-SNAP (http://www.itksnap.org/). A mesh of each participant's cortical surface was generated from the boundary of the white and gray matter.

***Delineation of primary V1, CoS-places, and mFus-faces.*** Regions of interest (ROIs) were delineated using cortex-based alignment of existing adult FreeSurfer brain atlases projected onto cortical surfaces of each infant, child, and adult participant at each time point. Specifically, the adult Wang atlas (*100*) was used to define primary visual cortex (V1) and the Rosenke atlas (*83*) for CoS-places and mFus-faces. The accuracy of the delineated ROIs was validated by comparing the automatically aligned brain structures to manually defined landmarks, such as the calcarine sulcus (Calc), using the Desikan atlas in prior work (*46*) and matched the dice coefficient from adult to adult brain.

***Analysis of $R_1$ and its development in cortical regions.*** To quantify developmental changes in cortical regions, -$R_1$ metrics were calculated for each ROI at each time point. Linear mixed models (LMMs) were employed to analyze the relationship between $R_1$ vs age and ROI age and metrics for each cortical region, incorporating random intercepts and slopes to account for individual variability.

**Comparison of qMRI and IHC in postmortem samples:**
***Postmortem MRI acquisition.*** Samples were placed in large plastic containers filled with Fomblin, a non-protonated fluid. The tissue samples were matched in size to their respective containers to ensure stability during the scan. Fomblin was degassed for 12 hours at room temperature to remove any air bubbles and to equilibrate the samples to room temperature before imaging. $R_1$, $T_1$w, and $T_2$w were acquired from each human brain sample (**Supplementary Fig. S2**) at the CNI 3T scanner for the 5-month, 9-year, and 25-year samples at 0.5mm$^3$ resolution and at 7T (NexGen 7T Siemens scanner at UC Berkeley) at higher resolution (0.33 mm$^3$) for the 1.5-month-old. MRI sequence parameters for postmortem scans are listed in **Supplementary Table 3.**

***Co-registration of (MRI slices and immunostained cryosections in postmortem samples.*** After the scan, brains were blocked as described above, with photographs of anatomical annotations at each iterative cut acquired to retain sample location relative to the initial scanned hemisphere. MR slices were prescribed parallel to the sample surface to simplify the co-registration process. The human IHC dataset was aligned with the 500 µm T2-weighted scan by manually identifying common vessels and tissue boundaries as landmarks and registering accordingly.

**Statistical analyses**
All statistical analyses were performed in MATLAB (R2024b) unless otherwise noted.

***IHC***
At least three sections were analyzed per experiment per sample at regularly spaced intervals across our ROI of interest in line with other publications in the field). Data were analyzed at the level of tissue sections and, where applicable, repeated measurements within a section (e.g., depth-wise sampling) were retained and modeled using mixed-effects frameworks to account for non-independence. All statistical analyses are reported in **Supplemental Table S4**.

***Mixed-effects modeling framework***
For outcomes measured across multiple sections and/or repeated measures per biological sample, we used linear mixed-effects models fit by restricted maximum likelihood (REML). Random intercepts were included to account for clustering within the relevant sampling unit (e.g., tissue section, slide, or donor/sample ID). Fixed effects included age (either categorical age group or continuous age, depending on the analysis) and anatomical region, and—when appropriate—their interaction. All tests were two-sided with $\alpha = 0.05$ unless stated otherwise.

***Myelin and MAP2 coverage across age and region***
To test developmental and regional effects in histological coverage measures (e.g., MBP/myelin coverage or MAP2 coverage), we fit REML mixed-effects models with fixed factors for Age Group and Region (and their interaction) and a random intercept for section. For example, myelin coverage was modeled with a full factorial specification: Coverage ~ Age_Group * Region + (1 | Section).

Model significance for fixed effects was assessed with F-tests (marginal ANOVA). Where a significant main effect and/or interaction was detected, post-hoc pairwise comparisons were conducted using two-sample Student's t-tests. Reported degrees of freedom and F-statistics correspond to the mixed-model ANOVA output, and model fit was summarized using the reported $R^2$ where applicable.

***$R_1$ analyses across age and region.***

$R_1$ was analyzed using linear mixed-effects models with a random intercept for subject to account for repeated regional measurements within an individual. Age was modeled as a continuous predictor using the log-transformed age variable to capture nonlinear developmental scaling, with region-by-age interaction: $R_1$ ~ Region * log(Age_days) + (1 | Subject). Models were fit for (i) the full dataset and (ii) an infant-restricted subset (defined as age $\leq$24 months). Region-specific slopes were derived from the fitted interaction terms, and planned pairwise contrasts of intercepts and slopes were evaluated using two-sided t-tests; effect sizes for planned comparisons were summarized using Cohen's d.

***Depth-wise relationships between $R_1$ and histology (MBP and MAP2)***

To quantify associations between MRI and histological measures across cortical depth while accounting for repeated measurements within samples, we used fixed linear mixed-effects models with random intercepts for sample ID: $R_1$ ~ MBP + (1 | ID) (across samples), $R_1$ ~ MBP + (1 | ID) and separately R1 ~ MAP2 + (1 | ID) (infant subset and child and adult subset). Model coefficients ($\beta$), standard errors, t-statistics, 95% confidence intervals, and p-values were reported. Model fit was summarized using AIC/BIC and log-likelihood.

***Reporting***

For each analysis, we report the model specification, sample sizes (number of sections and/or subjects, and number of repeated measurements where relevant), test statistics (F with numerator/denominator degrees of freedom for ANOVA; t for fixed effects and planned contrasts), and corresponding p-values. Where multiple post-hoc comparisons were performed, results are reported explicitly and all other non-significant comparisons are noted.

**Data and code availability.**

Data and code related to all data analyses, figures, and statistical modeling is available on the VPNL GitHub: https://github.com/VPNL/bbmyelin.

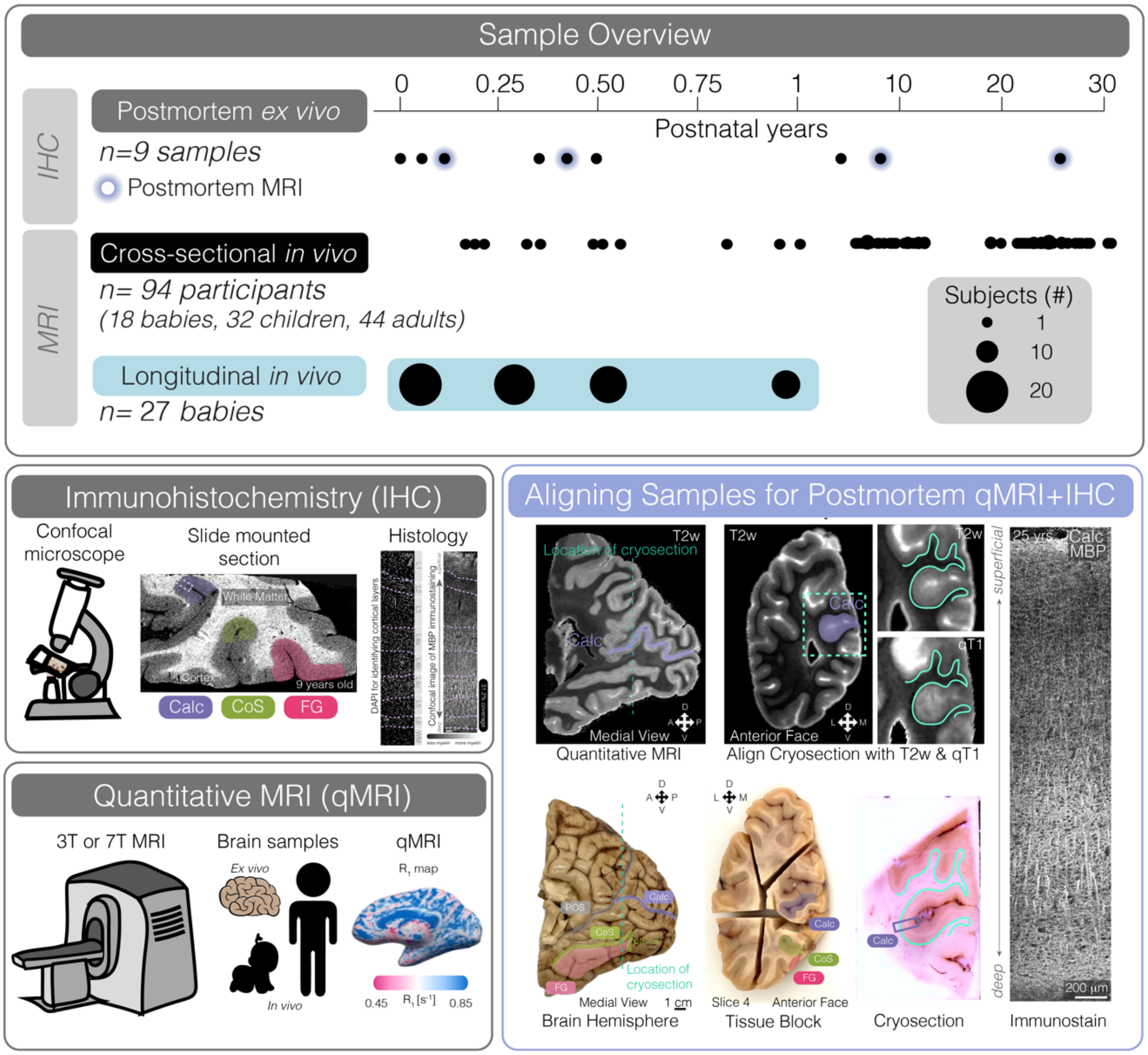


**Fig. S1. A comprehensive dataset of *in vivo* and postmortem visual cortex development.** Overview of pipeline for processing samples. *Top*: We conducted IHC on 9 postmortem samples, 4 of which were imaged with qMRI (blue cloud). 94 participants ages 0-30 years old were imaged cross-sectionally, and 27 babies were images longitudinally across their first year of life. *Bottom Left:* Pipeline for analyzing postmortem brains with immunohistochemistry (top) and quantitative MRI (qMRI) bottom in postmortem and *in vivo* samples. *Bottom right:* Method for aligning samples for postmortem qMRI and IHC. Hemispheres were first imaged with qMRI. Areas were identified in qMRI and on hemispheres. Tissue was then blocked into sections containing our areas of interest (bottom row, “Tissue Block”) and then cryosectioned (“Cryosection”) before immunostaining for MBP (“Immunostain”).

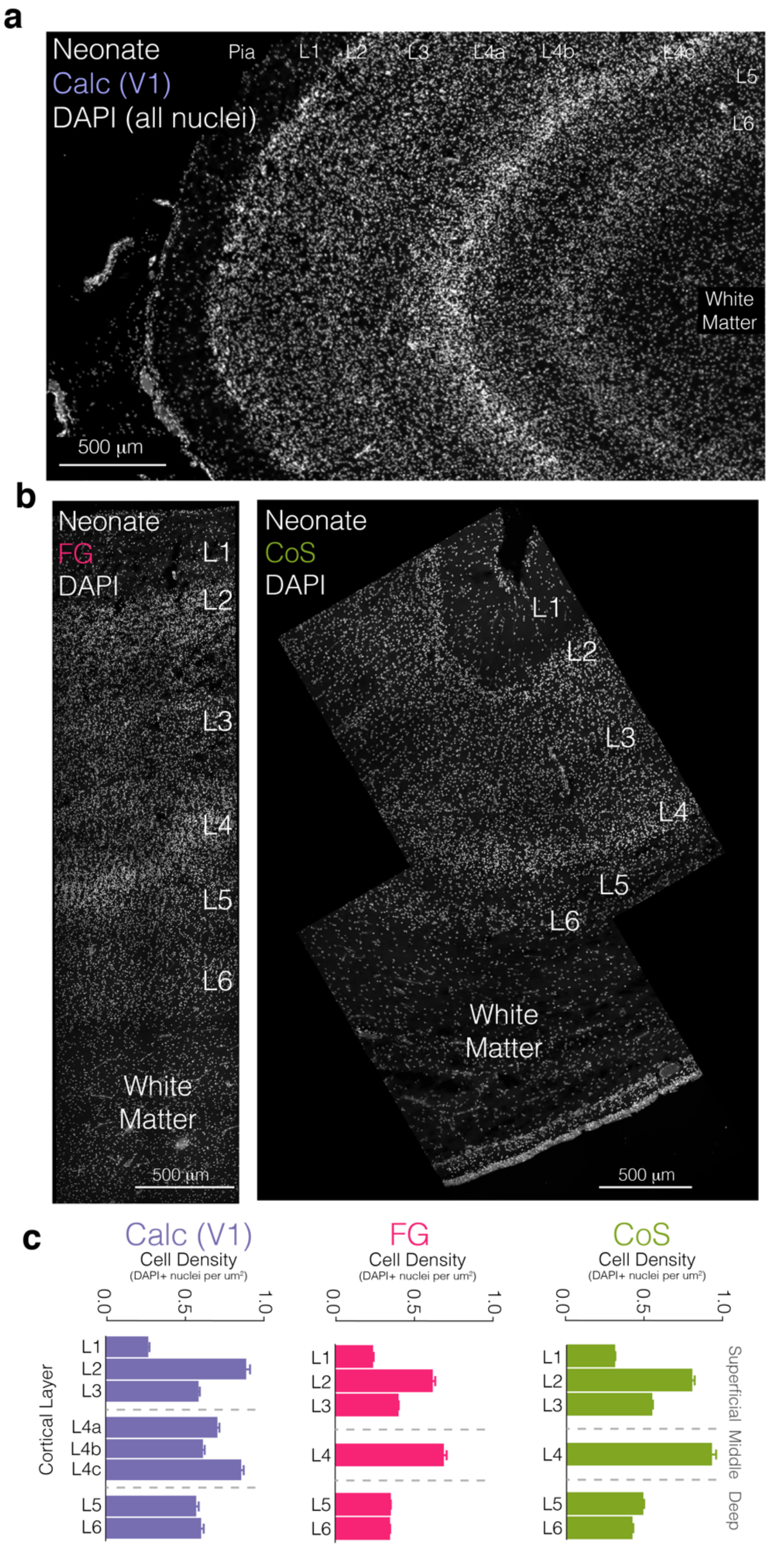


**Fig. S2. Identifying cortical layers with DAPI nuclear staining.**

**(a,b)** Cortical layers as identified by DAPI in Calc (**a**), FG (**b**, left), and CoS (**b**, right) in a neonatal brain sample (1mo). (**c**) Layers identified by changes in DAPI density.

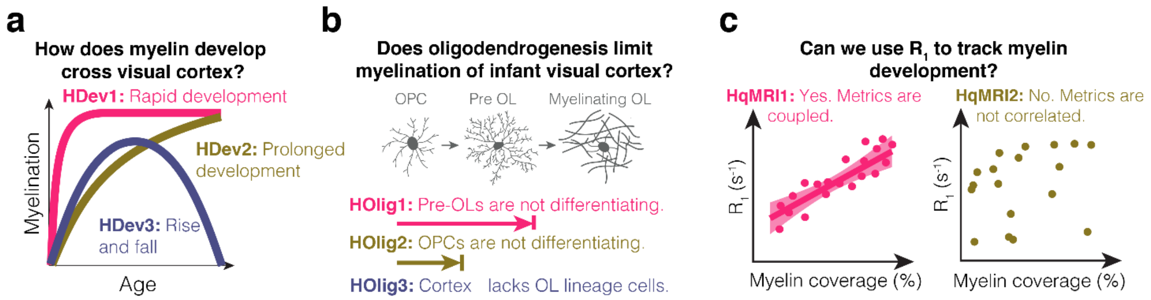


**Fig. S3. Proposed hypotheses,**

(**a**) **Hypothesized trajectories of myelin development**: early and rapidly (H1), prolonged across age (H2), or rise and then fall (H3). (**b**) **Hypotheses related to oligodendrogenesis:** In areas that lack myelin, oligodendrogenesis may be incomplete due to several reasons: (pre-OLs do not differentiate into myelinating OLs (H1, top), OPCs do not differentiate into pre-OLs (H2, middle), or developing cortex lacks oligodendrocyte lineage cells altogether (H3, bottom). (**c**) **Hypotheses related to relation between $R_1$ and myelin:** $R_1$ is modulated by myelin, iron, and tissue density (*66*, *71–73*, *90*). In adult visual cortex, higher $R_1$ is coupled with higher myelin, thus predicting that $R_1$ will increase with myelin across development (H1). Alternatively, the physiochemical environment of the cortex may be immature, hence the relationship between $R_1$ and myelin may not hold in sparsely myelinated infant cortex, where other microstructural factors may dominate $R_1$ (e.g., tissue density).

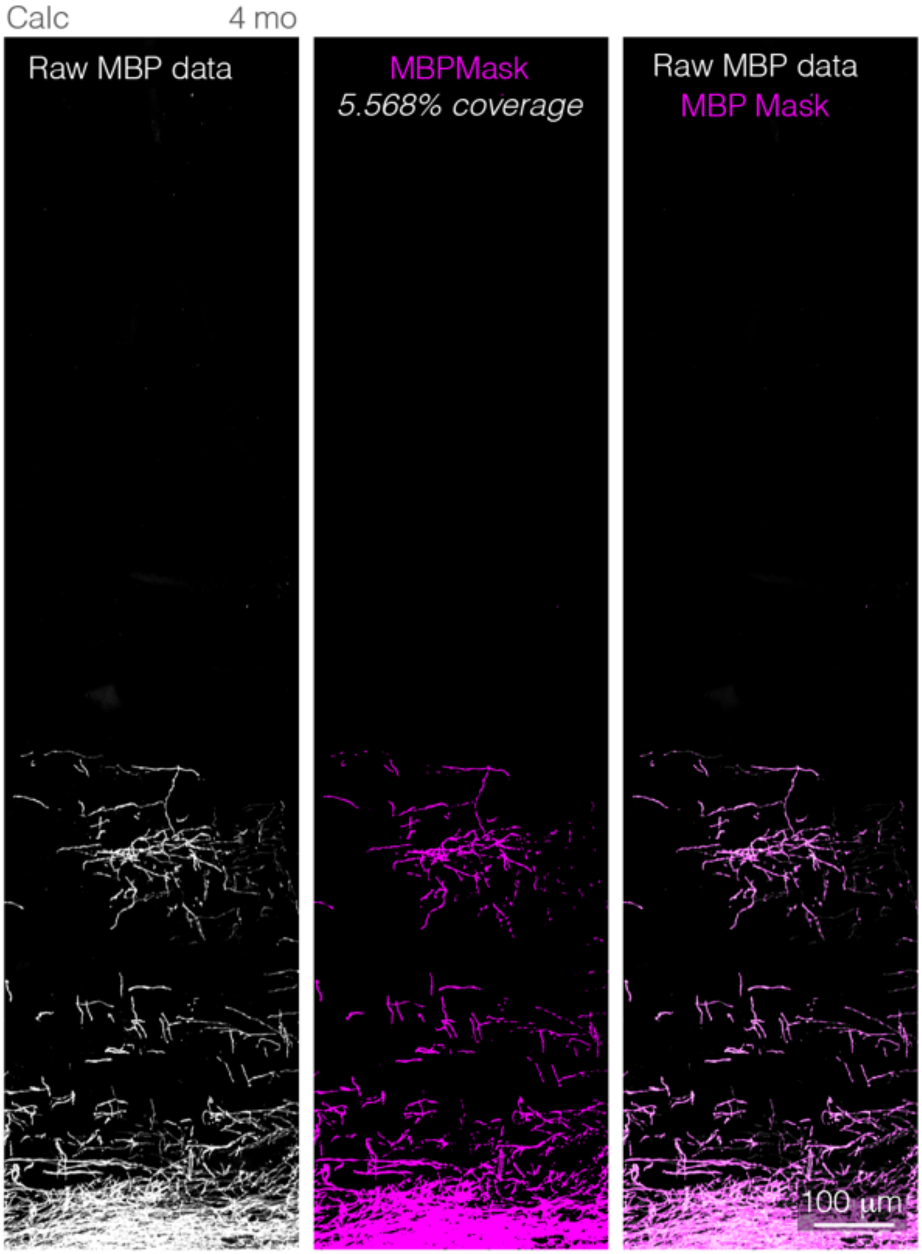


**Fig. S4. Quantifying myelin coverage using Otsu's automated thresholding method.**
Raw MBP data from max projections in a 4-month-old calcarine sulcus (Calc) sample (*left*) are processed using Otsu automated thresholding and then the total area covered by the MBP mask is computed using Analyze particles (5.568% coverage, in this case) (*middle*). Overlay of raw data with MBP mask shows high concordance between mask and raw data.

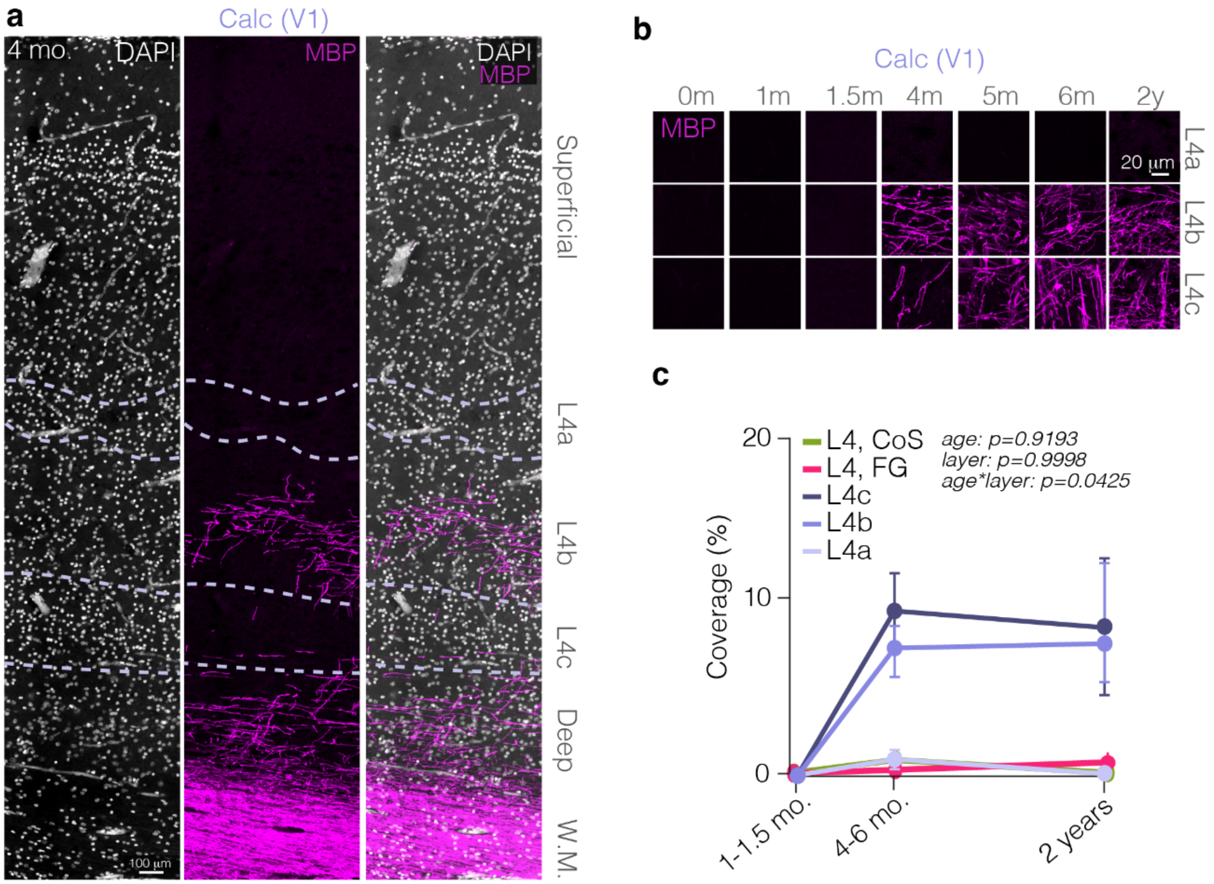


**Fig. S5. Myelin development in L4, including sublayers L4a-4c of Calc, of infant visual cortex.**

**(a)** Layer identification in a maximum projection of DAPI (left) and MBP (middle) immunostaining in a 4month old calcarine sulcus (Calc) sample. DAPI and MBP immunostain overlaid (right) to enable analysis of myelin coverage by layer. (**b)** Changes to myelin coverage in sublayers of L4 (L4a-c) in the calcarine sulcus. (**c**) Quantification of myelin coverage development in L4. In V1, L4b and L4c begin to myelinate in infancy, but L4a and L4 of CoS and FG remain myelin sparse throughout infancy.

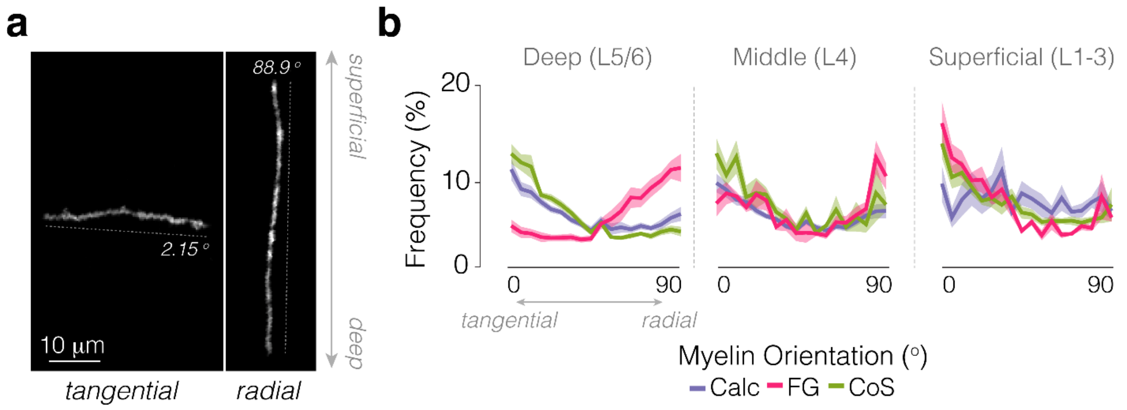


**Fig. S6. The orientation of deep layer myelination is area specific.**
**(a)** Maximum projection of an MBP-labeled myelin sheath oriented tangentially (left) and radially (right). **(b)** Myelin sheath orientation relative to the pial surface in deep (L5/6, left), middle (L4), and superficial (L1-3) cortex in calcarine sulcus (Calc), purple, fusiform gyrus (FG), pink, and collateral sulcus (CoS), green.

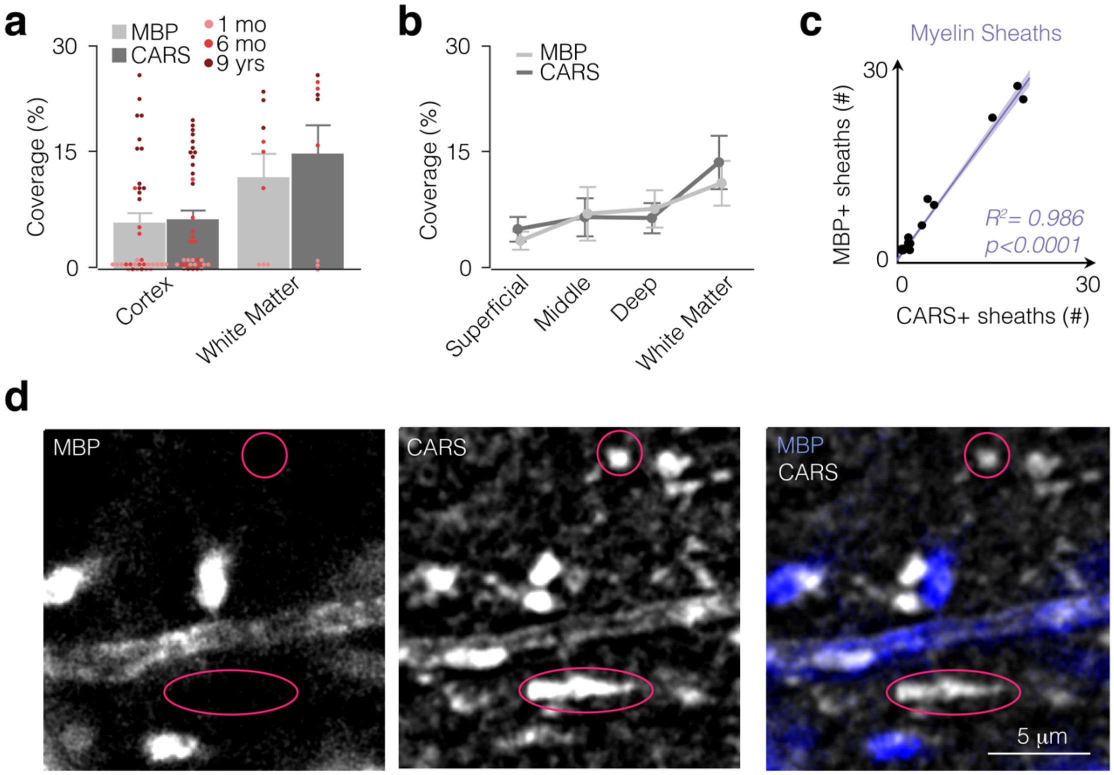


**Fig. S7. CARS (lipids) and MBP-immunostaining identify myelin sheaths in the cortex and superficial white matter underlying visual cortex.**
**(a)** MBP (light grey) and CARS (dark grey) identify similar levels of myelin coverage in co-imaged samples both in primary visual cortex in the calcarine sulcus (Calc) and the adjacent superficial white matter. **(b)** Patterns of MBP immunolabeling (light grey) and CARS (dark grey) coverage match across cortical depth. **(c)** In myelin sparse areas, the number of myelin sheaths identified with CARS and MBP-immunostaining is well-correlated, however, CARS identifies fewer myelin sheaths than MBP, likely due to myelin sheaths with low numbers of wraps, generating a low signal which may be undetectable in the CH-dense tissue matrix. **(d)** Max projection of a cortical section in calcarine sulcus (Calc, 6-month-old, L6) of MBP and CARS co-imaging. Pink circles demarcate examples of lipid features identified with CARS that are not associated with MBP-labeling. These features may be myelin/cell debris, intracellular or lysosome-associated lipid droplets, or compact, lipid-rich features, which can be investigated in future research. Thus, label-free CARS imaging reveals lipid features in developing human cortex, even before MBP expression, and validates our finding that the primary visual cortex (V1) does not contain mature myelin at birth.

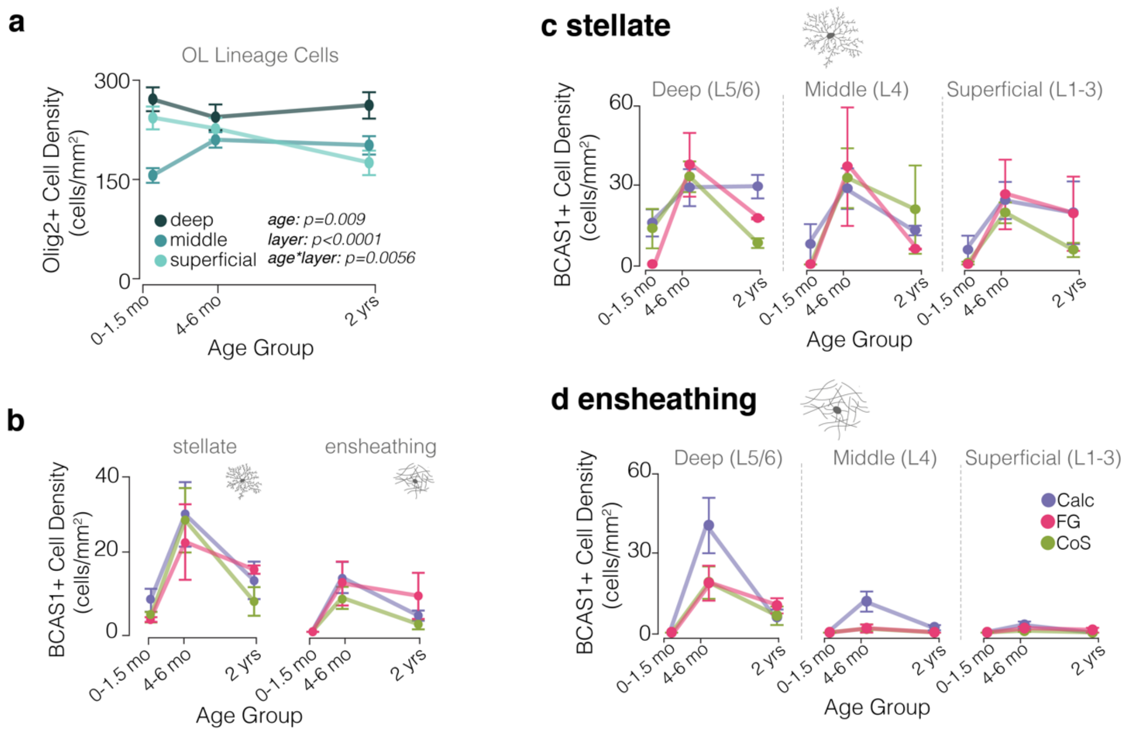


**Fig. S8. Oligodendrocyte lineage cells show similar patterns of development across visual areas.**
**(a)** Olig2+ cell density in deep (dark blue), middle (teal), and superficial (light teal) across cortex from 0-2 years old. **(b)** BCAS1+ cell density in Calc (purple), FG (pink), and CoS (green) across cortex from 0-2 years old. **(c)** BCAS1+ stellate (left, putative preOLs) and ensheathing (right, putative new OLs) cell densities in Calc (purple), FG (pink), and CoS (green) across cortex from 0-2 years old. **(d,e)** Density of BCAS1+ stellate (**d**) and ensheathing (**e**) cells in deep (left), middle (middle), and superficial (right) layers of cortex in Calc (purple), FG (pink), and CoS (green) across cortex from 0 to 2 years old.

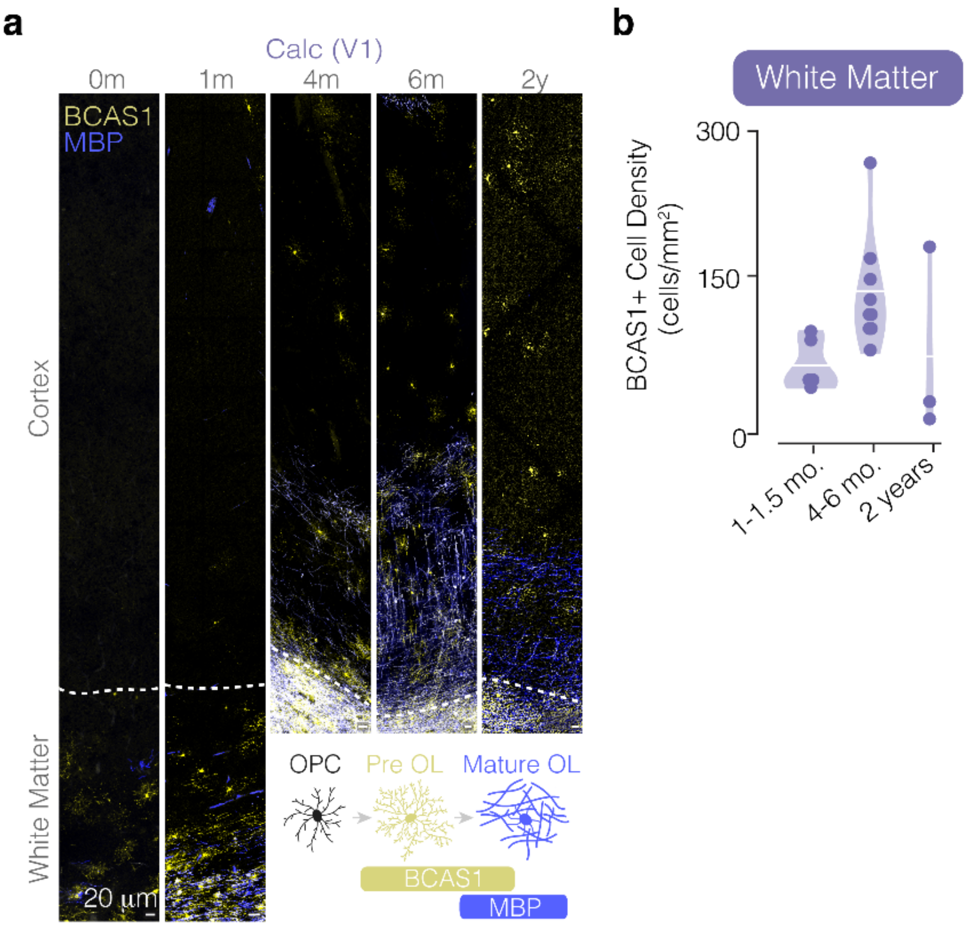


**Fig. S9. BCAS1+ cells are present at birth in the superficial white matter.**
**(a)** Maximum projection of MBP (blue) and BCAS1+ (yellow) staining in Calcarine sulcus (Calc) in a 0, 1, 4, and 6-month-old, and 2-year-old. Inlay shows development and time course of BCAS1 and MBP gene expression from OPCs into preOLs (labeled with BCAS1) and then mature OLs (labeled with MBP). **(b)** Density of BCAS1+ cells in the white matter adjacent to the calcarine sulcus from 0 to 2-years-old.

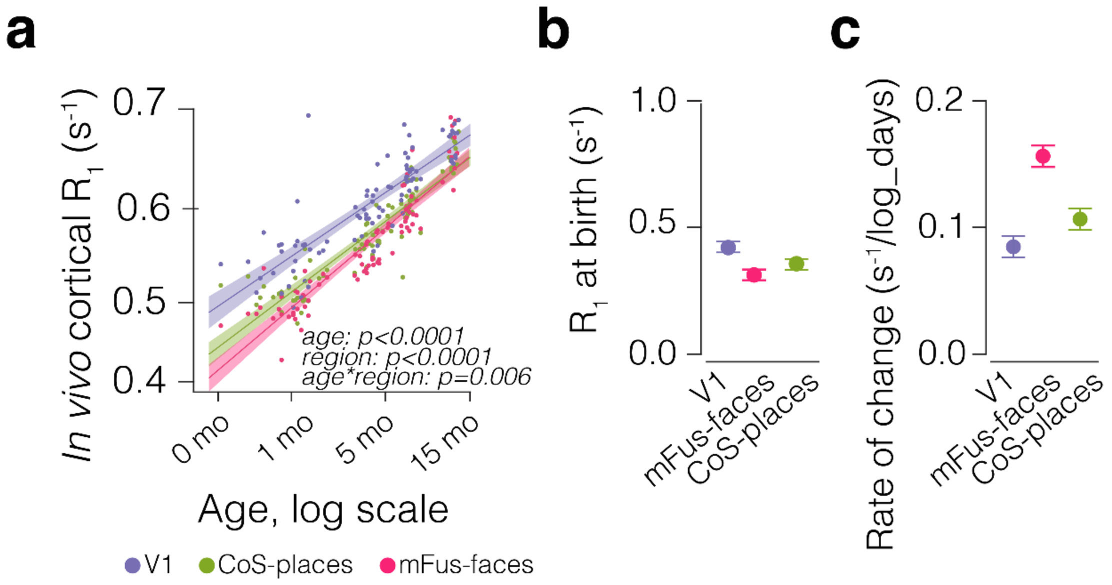


**Fig. S10. Changes to $R_1$ across development.**
**(a)** Longitudinal, *in vivo* cortical $R_1$ by log of age. **(b)** R1 at birth in Calc (purple), mFus-faces (pink), and CoS-places (green), calculated as the intercept of the line of fit in (a) for each area. **(c)** Rate of change of *in vivo* cortical $R_1$ from birth to 15 months of age, calculated as the slope of the line of fit in (a) for Calc (purple), mFus-faces (pink), and CoS-places (green).

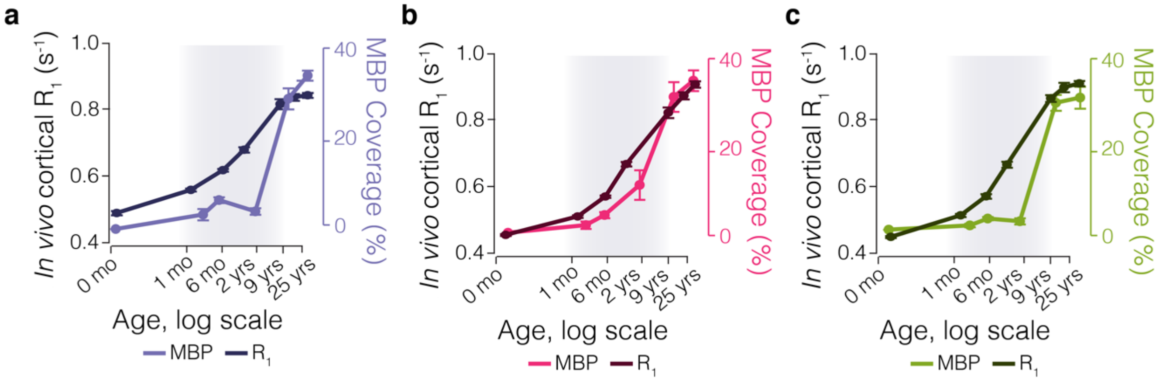


**Fig. S11. $R_1$ and MBP coverage across age in each region.**
**(a-c)** $R_1$ (darker color) and MBP coverage (lighter color) in Calc/V1 (purple, **a**), FG/mFus-faces (pink, **b**), and CoS-places (green, **c**) from 0-25 years. *Dots and error bars*: mean and SEM across scans ($R_1$)/sections (MBP). *Shaded gray area:* zone of discrepancy.

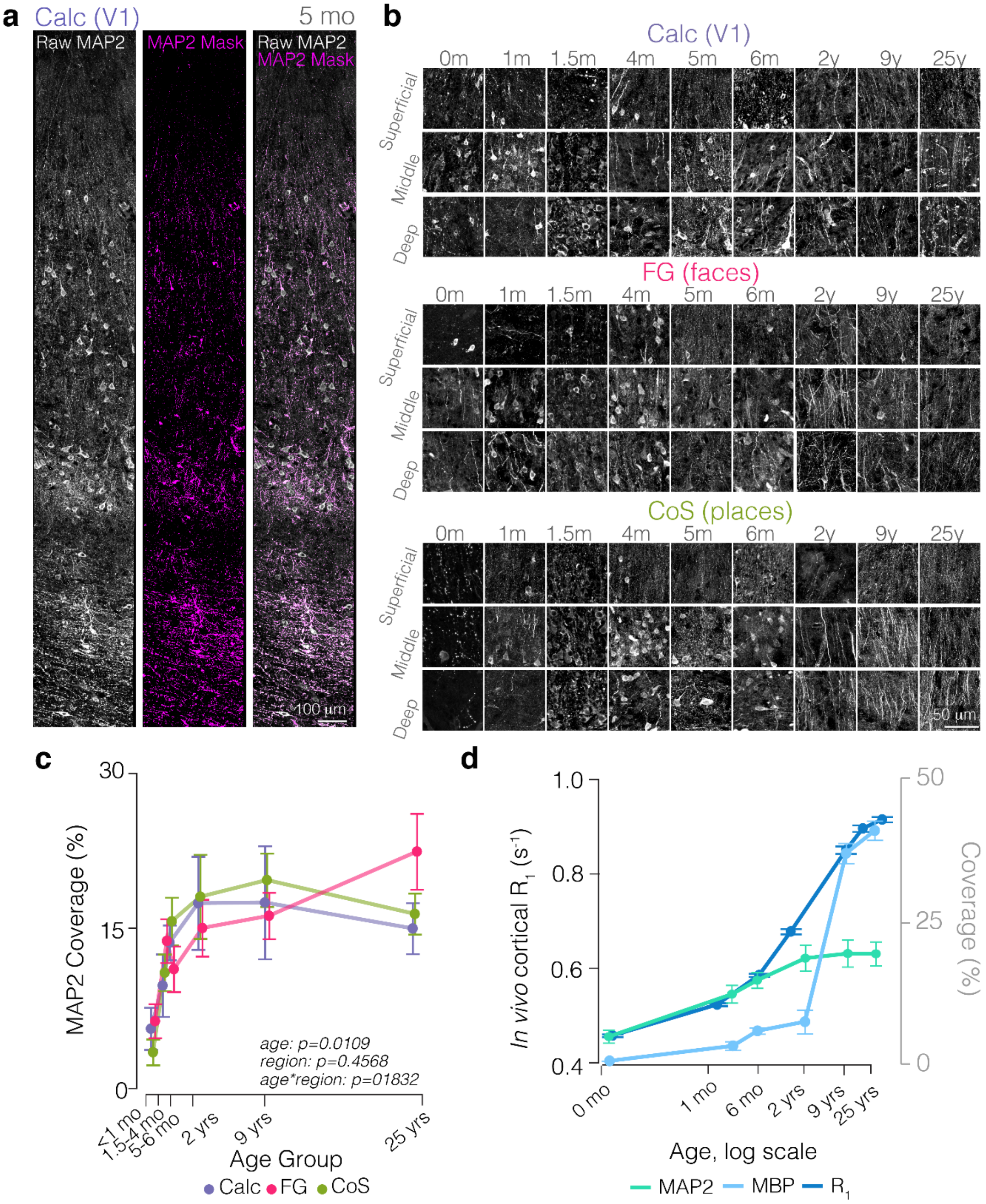


**Fig. S12. MAP2 develops rapidly from 0-4 months.**
**(a)** A maximum projection of MAP2 immunostaining (white, left) , a mask of MAP2 staining with cell bodies removed from the mask (pink, middle), and the overlay (right) of the raw data (white) and mask (pink), in a 5-month-old Calc sample. **(b)** Maximum projections of MAP2 immunostaining across cortical layers (superficial, top; middle, middle; deep, bottom) in 0-25 year olds in Calc (purple), FG (pink), and CoS (green). (**c**) MAP2 coverage in newborns (<1 month old), 1.5-4-month-olds, 5-6-month-olds, a 2-year-old, a 9-year-old, and a 25-year-old. This shows the increase in MAP2 that occurs between 0-4 months of age, in line with increases seen in $R_1$. **(d)** MAP2 (green), MBP (light blue), and *in vivo* $R_1$ (dark blue) development as a

function of age. $R_1$ aligns well with MAP2 growth in infants, then MAP2 plateaus and $R_1$ continues to grow, matching MBP by 9 years of age.

**Table S1. Supplementary Table 1 | Clinical and experimental demographics of collected human specimens.**

| Postnatal Age | Sex | Hemisphere | Clinical History | Experiments |
|---|---|---|---|---|
| 0 mo (1 postnatal day, born at 39 Gestational weeks. | F | L | Diaphragmatic repair | IHC |
| 1 mo (born at 35 GW) | M | Left | Pentalogy of Cantrell, low cardiac output state | IHC |
| 1.5 mo | M | Right | Respiratory distress Cardiomyopathy and respiratory distress | IHC + postmortem MRI |
| 4 mo | F | Left | Diaphragmatic repair | IHC |
| 5 mo | M | Left & Right | Congenital heart disease | IHC (LH, RH) + postmortem MRI (RH) |
| 6mo | M | Left | Cause of death was heart failure. Showed symptoms of seizures at 3 days old, symptoms resolved by 2 months. | IHC |
| 2 years | M | Left | Hemophagocytic lymphohistiocytosis, biliary tract obstruction and cholangitis | IHC |
| 9 years | F | Left & Right | Female, PAMI Syndrome, Subarachnoid and subdural hemorrhages, Disseminated intravascular coagulation | IHC (LH, RH) + postmortem MRI (LH) |
| 25 years | M | Right | Chronic myeloid leukemia and testicular cancer, diffused alveolar damage | IHC + postmortem MRI |

**Table S2. Supplementary Table 2 | Primary and secondary antibodies.**

| | Antigen | Host | Antibody Source | Catalog # | Dilution | RRID |
|---|---|---|---|---|---|---|
| Primary Antibodies | BCAS1 (NaBC1) | Mouse | Santa Cruz Biotechnology | sc-136342 | 1:200 | AB_10839529 |
| | MAP2 | Chicken | Abcam | ab5392 | 1:200 | AB_2138153 |
| | MBP | Rat | EMD Millipore | MAB386 | 1:200 | AB_94975 |
| | NeuN | Rabbit | EMD Millipore | ABN91 | 1:200 | AB_11205760 |
| | Olig2 | Rabbit | EMD Millipore | AB9610 | 1:100 | AB_570666 |
| Secondary Antibodies | Alexa Fluor 488-conjugated anti-chicken IgG | Donkey | Jackson | 703-545-155 | 1:200 | AB_2340375 |
| | Alexa Fluor 594-conjugated anti-rabbit IgG | Donkey | Invitrogen | A21207 | 1:200 | AB_141637 |
| | Alexa Fluor 594-conjugated anti-rat IgG | Donkey | Invitrogen | A21209 | 1:200 | AB_2535795 |
| | Alexa Fluor 647-conjugated anti-mouse IgG | Goat | Abcam | 150115 | 1:200 | AB_2687948 |
| | Alexa Fluor 647-conjugated anti-rat IgG | Goat | Invitrogen | A11007 | 1:200 | AB_141374 |

**Supplementary Table 3 | MRI Parameters.**

| MRI | Scan Type | Sample | Parameters | Length | Outcomes |
|---|---|---|---|---|---|
| aMRI | **Whole brain anatomy** | *in vivo* | BRAVO pulse sequence; resolution= 1mm^3, inversion time=450ms, TE=2.912 ms, FA=12 degrees | 3 min 15 sec | Whole brain volume & cortical surface reconstruction |
| | | *postmortem* | BRAVO pulse sequence; resolution= 0.5mm^3, inversion time=450ms, TE=3.62 ms, FA=11 degrees | 19 min 52 sec | |
| qMRI | **ViSTa-MRF: visualization of short transverse relaxation time with MR fingerprinting** | *in vivo* | 2D fast gradient Echo, FOV=22x22x22cm, resolution=2mm^3, flip angle: 90 degrees, receiver bandwidth = 125 Hz, # echoes = 1, TE = 1.5ms, TR = 12.5ms, with and without MT | 6 min 50 sec | Quantitative metrics of $R_1$ [$s^{-1}$], proton density (PD), myelin water fraction (MWF) |
| | | *postmortem* | 2D fast gradient Echo, FOV=22x22x22cm, resolution=0.5mm^3, flip angle: 90 degrees, receiver bandwidth = 125 Hz, # echoes = 1, TE = 3ms, TR = 12.5ms, with and without MT | 27 min 58 sec | |

**Supplementary Table 4 | Statistical Analyses.**

| Figure 1 | Measure | Statistical test |
|---|---|---|
| Fig. 1b | Myelin coverage (%) of individual sections in Calc, CoS, and FG across age groups (0-1.5 mo, 4-6 months, and 2 years) | Restricted Maximum Likelihood model (REML) to predict myelin coverage with full factorial model between Age Group (0-1.5mo vs. 4-6mo vs. 2yrs) and Region (Calc vs. FG vs. CoS), and with random variable of section.<br>Main effect of Age Group to predict coverage: $F(2,61)=23.14$, $p<0.0001$<br>Main effect of Region to predict coverage: $F(2,61)=0.02$, $p=0.9786$<br>Interaction effect between Age Group and Region to predict coverage: $F(4,61)=10.11$, $p<0.0001$<br>R2=0.79 |
| Fig. 1c | Myelin coverage (%) of individual sections in Calc, CoS, and FG across age groups (0-1.5 mo, 4-6 months, 2 years, 9 years, and 25 years) | Restricted Maximum Likelihood model (REML) to predict myelin coverage with full factorial model between Age Group (0-1.5mo vs. 4-6mo vs. 2yrs vs. 9yrs vs. 25yrs) and Region (Calc vs. FG vs. CoS), and with random variable of section.<br>Main effect of Age Group to predict coverage: $F(4,81)=140.45$, $p<0.0001$<br>Main effect of Region to predict coverage: $F(2,81)=0.01$, $p=0.9881$<br>Interaction effect between Age Group and Region to predict coverage: $F(8,81)=3.51$, $p=0.0016$<br>R2=0.98 |

| | | |
|---|---|---|
| Fig. 1d | Myelin coverage (%) of individual sections in Calc, CoS, and FG across age groups (0-1.5 mo, 4-6 months, 2 years, 9 years, and 25 years) and layer (deep, middle, and superficial) | Restricted Maximum Likelihood model (REML) to predict myelin coverage with full factorial model between Age Group (0-1.5mo vs. 4-6mo vs. 2yrs vs. 9yrs vs. 25yrs), Region (Calc vs. FG vs. CoS), and Layer (deep vs. middle vs. superficial) with random variable of section.<br>Main effect of Age Group to predict coverage: $F(4,249)=38.03$, $p<0.0001$<br>Main effect of Region to predict coverage: $F(2,249)=0.00$, $p=0.9979$<br>Main effect of Layer to predict coverage: $F(2,249)=0.00$, $p=0.9998$<br>Interaction effect of Age Group and Region to predict coverage: $F(8,249)=4.08$, $p<0.0001$<br>Interaction effect of Age Group and Layer to predict coverage: $F(8,249)=13.13$, $p<0.0001$<br>Interaction effect between Region and Layer to predict coverage: $F(4,249)=0.00$, $p=1.0000$<br>Interaction effect between Region. Layer, and Age Group to predict coverage: $F(16,249)=1.70$, $p=0.0467$<br>$R2=0.92$ |
| Fig. 1g | Relationship between the CARS-labeled myelin coverage and MBP-labeled myelin coverage | Standard least squares to predict change in CARS coverage using MBP coverage<br>Model: CARS = 1.2710 + 0.9446 * MBP<br>Intercept: 1.2710 (SE = 0.7003, p = 7.650413e-02)<br>Slope: 0.9446 (SE = 0.0630, p = 1.101408e-18)<br>$R^2$: 0.8394<br>Adjusted $R^2$: 0.8357<br>RMSE: 3.5054 |
| **Supplemental Fig. S5** | **Measure** | **Statistical test** |
| Fig. S5c | Myelin coverage (%) of individual sections in Calc L4 sublayers (4a, 4b, and 4c) and L4 of CoS and FG across age groups (0-1.5 mo, 4-6 months, and 2 years) | Restricted Maximum Likelihood model (REML) to predict myelin coverage with full factorial model between Age Group (0-1.5mo vs. 4-6mo vs. 2yrs) and Layer (4a vs. 4b vs. 4c), and with random variable of section.<br>Main effect of Age_group to predict coverage: $F(2,78)=0.09$, $p=0.9117$<br>Main effect of Layer to predict coverage: $F(4,78)=0.01$, $p=0.9998$<br>Interaction effect between Age_group and Layer to predict coverage: $F(8,78)=2.78$, $p=0.0094$<br>$R^2=0.74$ |
| **Supplemental Fig. S6** | **Measure** | **Statistical test** |

| | | |
|---|---|---|
| Fig. S6b | Myelin orientation (in degrees) of individual sheaths in Calc, CoS, and FG across cortical layers (deep, middle, and superficial) | Restricted Maximum Likelihood model (REML) to predict myelin orientation with full factorial model between Layer (Deep vs. Middle vs. Superficial) and Region (Calc vs. FG vs. CoS), and with random variable of section.<br>Main effect of Layer: $F(2,28)=0.11$, $p=0.899390$<br>Main effect of Region: $F(2,28)=2.16$, $p=0.133614$<br>Interaction effect between Layer and Region: $F(4,28)=7.09$, $p=0.000450$<br>$R^2=0.6735$ |
| **Supplemental Fig. S7** | **Measure** | **Statistical test** |
| Fig. S7a | Myelin coverage based on measure (CARS vs. MBP) across brain area (Cortex vs. WM) | Restricted Maximum Likelihood model (REML) to predict Coverage with full factorial model between Measure (CARS vs. MBP) and Brain Area (Cortex vs WM), and with random variable of section.<br><br>Main effect of Measure: $F(1,86)=0.03$, $p=0.8537$<br>Main effect of Brain Area: $F(1,86)=8.03$, $p=0.0057$<br>Interaction effect between Measure and Brain Area: $F(1,86)=0.36$, $p=0.5512$<br>$R^2=0.61$ |
| Fig. S7b | Myelin coverage based on measure (CARS vs. MBP) across depth (L1 vs. L2 vs. L4 vs. L6 vs. WM) | Main effect of Measure: $F(1,80)=0.18$, $p=0.6732$<br>Main effect of Layer: $F(4,80)=2.03$, $p=0.0986$<br>Interaction effect between Measure and Layer: $F(4,80)=0.19$, $p=0.9410$<br>$R^2=0.63$ |
| Fig. S7c | Relationship between the CARS-labeled myelin sheath number and MBP-labeled myelin sheath number | Standard least squares to predict change CARS sheat number using MBP sheath number<br>R-square = 0.99, $p<0.0001$ |
| **Figure 2** | **Measure** | **Statistical test** |

| | | |
|---|---|---|
| One-sample t-test to measure whether Olig2 density is greater than 0 at birth | One-sample t-test to measure whether Olig2 density is greater than 0 at birth | Calc - Deep:<br>t(200) = 8.7971<br>Calc - Middle:<br>t(536) = 11.8685<br>Calc - Superficial:<br>t(266) = 9.2489<br>CoS - Deep:<br>t(218) = 9.7735<br>CoS - Middle:<br>t(317) = 10.1826<br>CoS - Superficial:<br>t(355) = 12.2824<br>FG - Deep:<br>t(220) = 11.5273<br>FG - Middle:<br>t(160) = 8.9569<br>FG - Superficial:<br>t(87) = 6.4713 |
| Fig. 2e | Olig2 density (cells/mm^2) of individual sections in Calc, CoS, and FG across age groups (0-1.5 mo, 4-6 months, and 2 years) | Restricted Maximum Likelihood model (REML) to predict Olig2 density with continuous Age (in months), Region (Calc vs. FG vs. CoS), and Layer, with full three-way interaction, and with random variable of section.<br><br>Main effect of Age (continuous): F(1,5039)=0.14, p=0.7070<br>Main effect of Region: F(2,5039)=0.67, p=0.5124<br>Main effect of Layer: F(2,5039)=8.37, p=0.0002<br>Region × Age interaction: F(2,5039)=0.67, p=0.5124<br>Region × Layer interaction: F(4,5039)=0.38, p=0.8207<br>Age × Layer interaction: F(2,5039)=1.17, p=0.3094<br>Region × Age × Layer interaction: F(4,5039)=0.09, p=0.9864<br>$R^2$=0.04 |
| Fig. 2f | BCAS1 density (cells/mm^2) of individual sections in Calc, CoS, and FG across age groups (0-1.5 mo, 4-6 months, and 2 years) | Restricted Maximum Likelihood model (REML) to predict BCAS1 density with full factorial model between Age Group (0-1.5mo vs. 4-6mo vs. 2yrs) and Region (Calc vs. FG vs. CoS), and with random variable of section.<br>Main effect of Age Group: F(2,39)=5.15, p=0.0104<br>Main effect of Region: F(2,39)=0.13, p=0.8810<br>Interaction effect between Age Group and Region: F(4,39)=0.29, p=0.8799 |

| | | |
|---|---|---|
| Fig. 2g | BCAS1 density by morphology (stellate and ensheathing) of individual layers (deep vs. middle. vs. superficial) across age groups (0-1.5 mo vs. 4-6 months vs. 2 years) | Restricted Maximum Likelihood model (REML) to predict BCAS1 density with full factorial model between Age Group (0-1.5mo vs. 4-6mo vs. 2yrs) and Layer (deep,middle,superficial), and with random variable of section.<br>STELLATE<br>Main effect of Age_group (categorical): $F(2,132)=2.72$, $p=0.0694$<br>Main effect of Layer: $F(2,132)=2.19$, $p=0.1165$<br>Layer × Age_group interaction: $F(4,132)=1.09$, $p=0.3644$<br>$R^2=0.42$<br>ENSHEATHING<br>Main effect of Age_group (categorical): $F(2,132)=53.99$, $p<0.0001$<br>Main effect of Layer: $F(2,132)=0.00$, $p=0.9992$<br>Layer × Age_group interaction: $F(4,132)=13.06$, $p<0.0001$<br>$R^2=0.61$ |
| **Supplemental Fig. S8** | **Measure** | **Statistical test** |
| Fig. S8a | Olig2 density (cells/mm^2) of individual sections in Calc, CoS, and FG across age groups (0-1.5 mo, 4-6 months, and 2 years) and layers (deep, middle, superficial) | Restricted Maximum Likelihood model (REML) to predict Olig2 density with continuous Age (in months), Region (Calc vs. FG vs. CoS), and Layer, with full three-way interaction, and with random variable of section.<br><br>Main effect of Age (continuous): $F(1,5039)=0.14$, $p=0.7070$<br>Main effect of Region: $F(2,5039)=0.67$, $p=0.5124$<br>Main effect of Layer: $F(2,5039)=8.37$, $p=0.0002$<br>Region × Age interaction: $F(2,5039)=0.67$, $p=0.5124$<br>Region × Layer interaction: $F(4,5039)=0.38$, $p=0.8207$<br>Age × Layer interaction: $F(2,5039)=1.17$, $p=0.3094$<br>Region × Age × Layer interaction: $F(4,5039)=0.09$, $p=0.9864$<br>$R^2=0.04$ |
| Fig. S8b | BCAS1 density (cells/mm^2) of individual regions (Calc vs. FG. vs. CoS) across age groups (0-1.5 mo vs. 4-6 months vs. 2 years) | Restricted Maximum Likelihood model (REML) to predict BCAS1 density with full factorial model between Age Group (0-1.5mo vs. 4-6mo vs. 2yrs) and Region (Calc, FG, CoS), and with random variable of section.<br>STELLATE<br>Main effect of Age (continuous): $F(1,42)=0.01$, $p=0.9060$<br>Main effect of Region: $F(2,42)=0.05$, $p=0.9501$<br>Region × Age interaction: $F(2,42)=0.14$, $p=0.8729$<br>$R^2=0.41$<br>ENSHEATHING<br>Main effect of Age (continuous): $F(1,42)=0.00$, $p=0.9681$<br>Main effect of Region: $F(2,42)=1.51$, $p=0.2329$<br>Region × Age interaction: $F(2,42)=0.05$, $p=0.9494$<br>$R^2=0.66$ |

| Fig. S8c | BCAS1 stellate cell density (cells/mm^2) across age groups (0-1.5 mo vs. 4-6 months vs. 2 years) and Regions (Calc vs. FG vs. CoS) and Layers (superficial, middle, deep) | Restricted Maximum Likelihood model (REML) to predict BCAS1 density with full factorial model between Age Group (0-1.5mo vs. 4-6mo vs. 2yrs) and Region (Calc vs. FG vs. CoS), and Layer (deep, middle, superficial) with random variable of section.<br>Main effect of Age (continuous): $F(1,123)=0.82$, $p=0.3664$<br>Main effect of Region: $F(2,123)=0.60$, $p=0.5511$<br>Main effect of Layer: $F(2,123)=1.79$, $p=0.1717$<br>Region × Age interaction: $F(2,123)=0.38$, $p=0.6825$<br>Region × Layer interaction: $F(4,123)=0.70$, $p=0.5959$<br>Age × Layer interaction: $F(2,123)=0.38$, $p=0.6817$<br>Region × Age × Layer interaction: $F(4,123)=0.81$, $p=0.5191$<br>$R^2=0.43$ |
|---|---|---|
| Fig. S8d | BCAS1 ensheathing cell density (cells/mm^2) across age groups (0-1.5 mo vs. 4-6 months vs. 2 years) and Regions (Calc vs. FG vs. CoS) and Layers (superficial, middle, deep) | Restricted Maximum Likelihood model (REML) to predict BCAS1 density with full factorial model between Age Group (0-1.5mo vs. 4-6mo vs. 2yrs) and Region (Calc vs. FG vs. CoS) and Layer (deep,middle,superficial), and with random variable of section.<br>Main effect of Age (continuous): $F(1,123)=0.96$, $p=0.3280$<br>Main effect of Region: $F(2,123)=1.18$, $p=0.3122$<br>Main effect of Layer: $F(2,123)=8.91$, $p=0.0002$<br>Region × Age interaction: $F(2,123)=0.21$, $p=0.8112$<br>Region × Layer interaction: $F(4,123)=0.14$, $p=0.9652$<br>Age × Layer interaction: $F(2,123)=0.41$, $p=0.6670$<br>Region × Age × Layer interaction: $F(4,123)=0.09$, $p=0.9863$<br>$R^2=0.50$ |
| **Supplemental Fig. S9** | **Measure** | **Statistical test** |
| Fig. S9b | BCAS1 cell density in superficial white matter of Calcarine sulcus across age (0-1.5 mo vs. 4-6 months vs. 2 years) | Null hypothesis: Mean cell density = 0<br>Alternative hypothesis: Mean cell density ≠ 0<br><br>Age Group n Mean SEM t-stat p-value Sig<br>------------------------------------------------------------------------------------<br>0-1.5mo 6 68.46 9.82 6.9706 0.000935 ***<br>4-6mo 9 141.61 18.57 7.6267 0.000061 ***<br>2yrs 3 79.00 54.46 1.4506 0.283969 ns |
| **Figure 3** | **Measure** | **Statistical test** |

| | | |
|---|---|---|
| Fig. 3b | R1 of individuals in V1, CoS-places, and mFus-faces across age (0-15 months) | Restricted Maximum Likelihood model (REML) to predict R1 with full factorial model between Age (0-15mo) and Region (V1vs. mFus-faces vs. CoS-places), and with random variable of individual.<br>Main effect of log(Age_days): $F(1,254)=362.60$, $p<0.0001$<br>Main effect of Region: $F(2,254)=15.88$, $p<0.0001$<br>Interaction effect between log(Age_days) and Region: $F(2,254)=5.24$, $p=0.0059$<br>$R^2=0.86$ |
| Fig. 3c | R1 of individuals in V1, CoS-places, and mFus-faces across age (0-25 years) | Restricted Maximum Likelihood model (REML) to predict R1 with full factorial model between Age (0-25yrs) and Region (V1vs. mFus-faces vs. CoS-places), and with random variable of individual.<br>Main effect of log(Age_days): $F(1,482)=943.64$, $p<0.0001$<br>Main effect of Region: $F(2,482)=40.27$, $p<0.0001$<br>Interaction effect between log(Age_days) and Region: $F(2,482)=20.94$, $p<0.0001$<br>$R^2=0.93$ |
| **Figure 4** | **Measure** | **Statistical test** |
| Fig. 4b | Relationship between R1 and MBP coverage across sampes (1.5 month, 5 month, 9 year, and 25 year) | Fixed linear mixed-effects model to predict change in R1 using MBP coverage with random intercept of sample (ID). Formula: R1 ~ MBP + (1\|ID) |
| Fig. 4c | Relationship between R1 and MBP coverage or MAP2 coverage across infant sampes (1.5 month, 5 month) | Fixed linear mixed-effects model to predict change in R1 using MBP coverage with random intercept of sample (ID). Formula: R1 ~ MBP or MAP2 + (1\|ID) |
| Fig. 4d | Relationship between the R1 and MBP coverage or MAP2 coverage | Fixed linear mixed-effects model to predict change in R1 using MBP coverage with rrandom intercept of sample (ID). Formula: R1 ~ MBP or MAP2 + (1\|ID) |

| | | |
|---|---|---|
| | across infant sampes (9 years, 25 years) | |
| **Supplemental Fig. S12** | **Measure** | **Statistical test** |
| Fig. S12d | MAP2 coverage (%) of individual sections in Calc, CoS, and FG across age groups (0-1 mo, 1.5-4mo, 5-6 months, 2 years, 9 years, and 25 years) | Restricted Maximum Likelihood model (REML) to predict MAP2 coverage with full factorial model between Age Group (0-1mo vs. 1.5-4mo vs. 5-6mo vs. 2yrs vs. 9yrs vs. 25yrs) and Region (Calc vs. FG vs. CoS), and with random variable of section.<br>Main effect of Age Group to predict coverage: $F(5,61)=3.28$, $p=0.0109$<br>Main effect of Region to predict coverage: $F(2,61)=0.79$, $p=0.4568$<br>Interaction effect between Age Group and Region to predict coverage: $F(10,61)=1.44$, $p=0.1832$<br>$R2=0.80$ |
